# Annotating Hypothetical Proteins in *Escherichia coli* K-12

**DOI:** 10.1101/2025.03.20.643886

**Authors:** Sagarika Chakraborty, Zachary Ardern, Habibu Aliyu, Anne-Kristin Kaster

## Abstract

Omics technologies have led to the discovery of a vast number of proteins that are expressed but have no functional annotation - so called hypothetical proteins (HPs). Even in the best-studied model organism *Escherichia coli* K-12, over 2% of the proteome remains uncharacterized. This knowledge gap becomes even worse when looking at microbial dark matter. However, knowing the functions of proteins is crucial for elucidating cellular and metabolic processes and harnessing biotechnological potentials. Here, we employed machine learning to decipher the transcriptional regulatory network of *E. coli* K-12, as well as other *in silico* tools to assign functions to uncharacterized HPs. We further provide experimental validation of *in silico* predicted functions for three HP-encoding genes (*yhdN*, *yeaC* and *ydgH*) as proof of concept, by analyzing growth patterns of deletion mutants compared to the wild type, as well as their transcriptional responses to specific conditions. This study demonstrates that the use of Big Omics Data in combination with Artificial Intelligence and experimental controls is a powerful approach to illuminate functional dark matter.

**Graphical Abstract:** 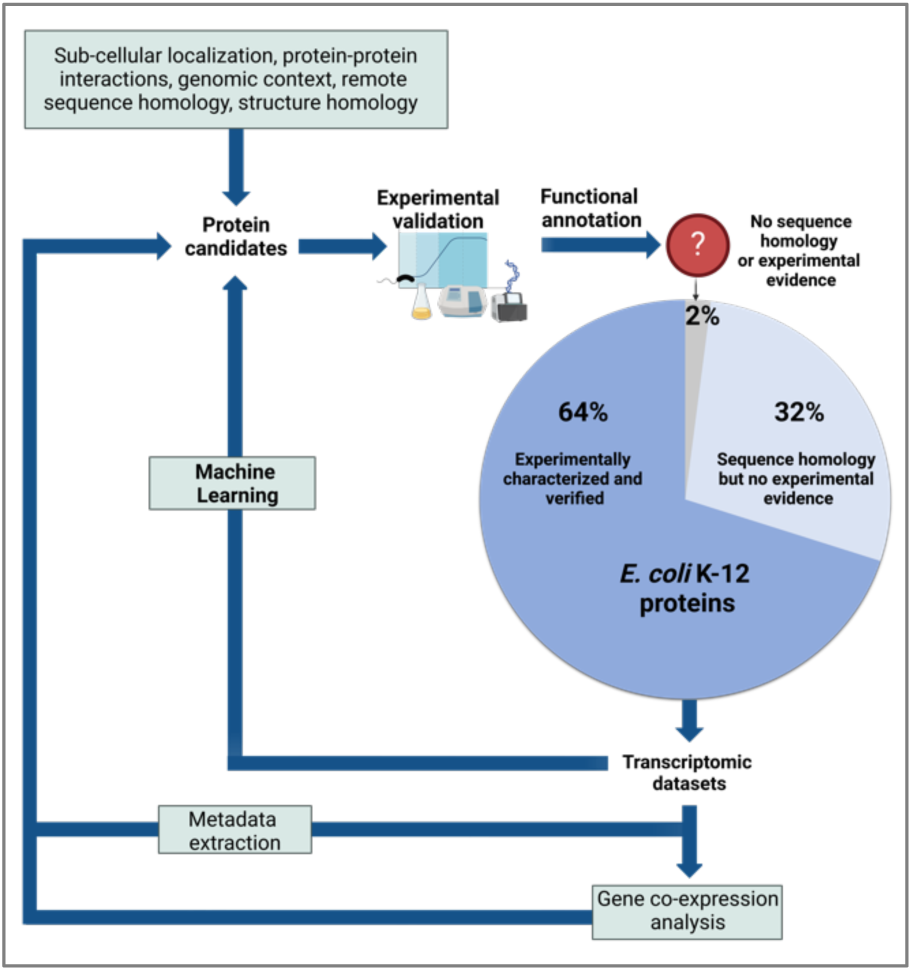

## Introduction

Due to the advents in Next Generation Sequencing technologies, the volume of omics data has massively increased over the past years. As of today, gold-standard knowledgebases like the National Center for Biotechnology Information (NCBI)’s Reference Sequences Database (RefSeq) and the Universal Protein Resource Database (UniProt) [1] provide freely accessible sequences and information for over 163 million unique prokaryotic proteins **(Figure 1)**. This rapid accumulation of sequencing data has long surpassed the rate of possible experimental characterization *in vivo* and *in vitro*. Hence, researchers often rely on functional analysis of proteins from model organisms. These findings are then extrapolated *in silico* to closely related sequence homologues. Consequently, the definition of cut-off values plays a crucial role in protein annotation. To manage this challenge, automated annotation pipelines, such as InterPro have become a valuable tool [2]. Here, analysis of protein sequences is provided by classifying them into families and integrating predictive models from multiple protein databases such as e.g. Pfam [3], ProSite [4], SMART [5] and/or CDD [6] to infer functions for experimentally uncharacterized proteins. However, only 83% of all UniProt sequences have so far been functionally annotated by InterPro [7] The remaining 17% of proteins are designated with terms such as ‘putative uncharacterized’, ‘uncharacterized’, ‘unknown protein families (UPFs)’ or proteins bearing ‘domains of unknown functions’ (DUFs). These proteins are summarized by the term hypothetical proteins (HPs), since they are expressed but could so far not be functionally characterized by classical *in vivo*, *in vitro*, and/or *in silico* methods (8,9). In addition, there is also no information on these proteins available in widely-used knowledge-databases such as BioCyc [10], RegulonDB [11], and EggNOG [12] **(Figure 1).**

**Figure 1.**
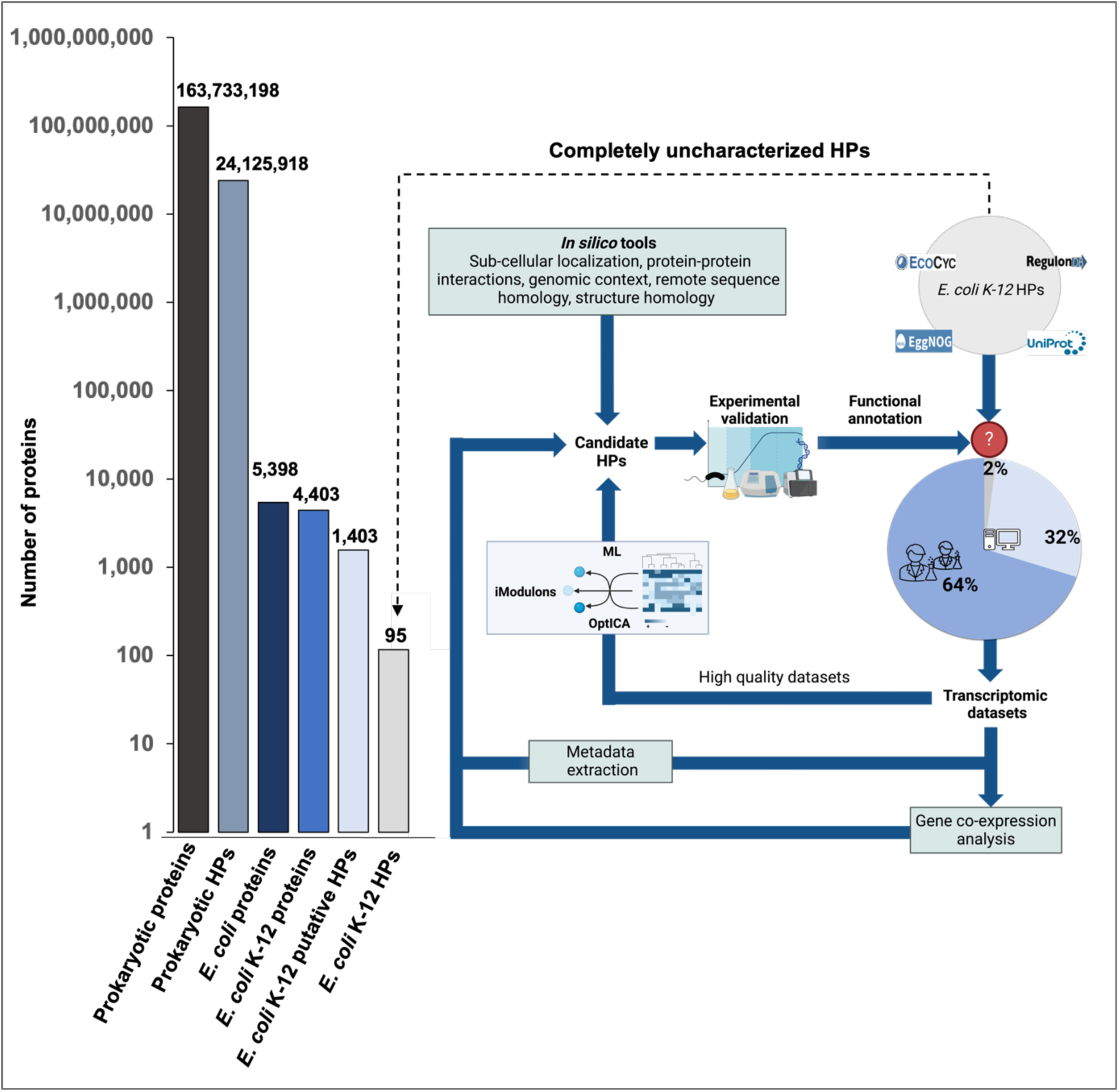
Total number of sequences for all unique prokaryotic and *Escherichia coli* proteins deposited in the National Center for Biotechnology Information (NCBI) as of April 2024 and methodological set-up of this study. Of the 4403 *E. coli* K-12 protein encoding genes, 1403 (31.8%) encode for unique proteins with functions predicted only *in silico* based on homologous sequences but lacking *in vivo* or *in vitro* experimental evidence (termed “putative hypothetical proteins”). 95 protein encoding genes (2.1%) of *E. coli* K-12 are completely uncharacterized with no sequence homologues according to the four knowledge databases - EcoCyc [39], RegulonDB [11], EggNOG [12] and UniProt (1) (termed “hypothetical proteins”). Transcriptomic datasets from NCBI were filtered and processed using the OptICA approach to generate iModulons [32]. Metadata information was curated in parallel using manual or semi-automated approaches(39–41).Bioinformatics, machine learning and deep learning tools along with the presence of relevant metadata then resulted in potential functions for HP candidates for *in vitro* testing. HPs, hypothetical proteins; ICA, independent component analysis; ML, machine learning.

While there have been notable advancements in methodologies for protein function prediction in the recent past [13–16] several challenges persist [17,18]: These include different functionalities across homologous proteins, proteins performing different functions in distinct cellular locations, proteins with multiple three-dimensional structures, the lack of conserved proteins across different species to identify shared functional relationships and a heavy reliance on pre-existing data for inferential annotations [19]. Even in the best studied model organisms *Escherichia coli* K-12, 31.8% of the genome comprises of protein encoding genes that lack experimental validation, commonly referred to as ‘Putative HPs’ [8], and 2.1% that even lack sequence homologies (this study) (**Figure 1**). The problem of annotating HPs becomes even larger when looking at microbial dark matter(9, 20).In uncultivated microorganisms, the proportions of genes with unknown functions can comprise up to 60% in bacterial and 80% in archaeal genomes [21]. However, elucidating the function of these proteins is essential for understanding cellular processes, metabolism and evolution, as well as for harnessing their biotechnological potential [22].

Transcriptional profiling has helped to gain insights on protein functions by monitoring gene expressions and clustering genes based on their responses to varying environmental conditions and stimuli [23]. Here, the entire set of RNA transcripts produced by a genome under specific conditions provides a dynamic representation of the organisms’ operational state. However, standard bioinformatics-based methods for transcriptomic data analysis are low-throughput and require quite extensive computing power and time due to the high degrees of complexity and heterogeneity of the data [24]. Today, artificial intelligence (AI)-based methodologies have made significant advancements over traditional bioinformatics approaches, offering the capacity to process and analyze data at a scale and speed that were previously unattainable [24, 25]. They are particularly suited for deciphering complex interactions among genes and transcription factors or proteins that regulate them, as well as revealing the regulation of gene expression in response to diverse environmental conditions [27]. Among various techniques for analysis, module-detection methods have proven to efficiently predict the functions of co-regulated groups of genes from large gene expression datasets [28]. Independent Component Analysis (ICA), introduced by Comon in 1994 [29], employs an unsupervised machine learning (ML) approach, categorizing the input data without any prior information. This technique is especially relevant in the context of transcriptomics, where the goal is to understand the complex interplay of gene expression signals and to decipher co-regulated gene sets. ICA therefore allows to reveal the regulatory patterns and activity levels of genes governed by specific regulators (i.e. transcription factors controlling the expression of genes) across diverse experimental conditions, so called transcriptional regulatory networks (TRNs), thereby disclosing the organizational and functional architecture of genes. Expanding upon this, McConn et al. developed a variant of ICA, termed OptICA, which can be applied to large TRNs, analyzing interactions between transcription factors and their target genes. Here, discrete groups of genes are clustered into robust independent components by avoiding over-clustering of datasets (leading to loss of biological information) or under-clustering (leading to too few clusters and a loss of output resolution), and therefore providing an optimal representation of the organism’s underlying TRN [27]. This results in revealing independently regulated gene groups, so called iModulon from transcriptomics datasets [29, 30]. iModulons are therefore the data-driven analogs of so-called regulons - sets of genes governed by regulators [30]. The OptICA algorithm along with high-quality RNA-seq data has recently been used to decipher the TRN of *Bacillus subtilis* [32] and *E. coli* [27] to uncover the detailed responses to environmental conditions and genetic perturbations, e.g. deletion mutants. However, these studies focused on the identification of regulons that were not experimentally described before rather than the functional characterization of HPs.

In order to decipher the function of HP-encoding genes in the *E. coli* K-12 proteome, we adapted the workflow originally designed for *Bacillus subtilis* [32] and analyzed the gene expression data of the *E. coli* K-12 sub-strains MG1655 and BW25113. Functional characterization of HP-encoding genes was inferred by identifying co-regulated genes under the influence of a common regulator from publicly available transcriptomic datasets. This characterization was then further validated by annotating these genes using specific Gene Ontology (GO) categories through the PANTHER tool (Protein ANalysis THrough Evolutionary Relationships) [33]. Additionally, metadata conditions for the transcriptomic datasets were extracted. Furthermore, other *in silico* methods were employed to determine the sub-cellular localization [34], protein-protein interactions (PPIs) [35], genomic context [36] and sequence-based remote homologies [37]. Structural homology information was obtained by using the AlphaFold Protein Structure Database (AFDB) clusters webserver, which is based on a deep learning (DL) method [38].

Since AI-predicted functions are usually not experimentally verified, raising the possibility of untrue annotations [9], we also provide an experimental validation of three candidate HP-encoding genes (*yhdN*, *yeaC* and *ydgH*). The growth curves and transcriptional responses of *E. coli* K-12 deletion mutants in comparison to the wild-type strain under specific conditions were analyzed, showing that the *in silico* predictions were indeed true. Hence, the here presented pipeline not only facilitates the discovery of previously unidentified regulators, but also sheds light on our understanding of the function of HPs **(Figure 1).**

## Results and Discussion

### Identification of uncharacterized HPs in *E. coli*

In 2019, Ghatak et al. had identified HP-encoding genes in *E. coli* K-12, however only by employing keywords such as ‘possibly’, ‘predicted’ or ‘hypothetical’ from the EcoCyc database, or annotation scores of two or below in UniProt [8]. This approach did, however, not establish a stringent threshold for accurate identification of completely uncharacterized HP-encoding genes. Hence, a more rigorous approach was used in this study, which included additional keywords for HPs (such as ‘Uncharacterized’, ‘Putative uncharacterized’, ‘DUF’ etc.) and information on the absence of any functional characterization using multiple gold-standard databases (EcoCyc, UniProt, RegulonDB and EggNOG). 158 of the 1600 HP-encoding genes previously listed by Ghatak et al. have now been functionally characterized with experimental evidence in the EcoCyc database. Notably, the HP-encoding gene *ydfX*, curated as one of the HPs in this study, was missing from Ghatak et al.’s list since it was previously categorized as a pseudogene/phantom gene. Its exclusion, however, could not be verified in the current EcoCyc database. We identified 1403 of 4403 genes (31.8%) of the *E. coli* K-12 genome as ‘putative’ HPs, implying that their functions were inferred based on sequence homologies in the gold-standard databases, but lacking an experimental evidence for function. A recent study using the EggNOG tool also identified about 30% in a different *E.* strain (O157:H7 strain Sakai) as putative HPs [18].

95 HP-encoding genes (2.1%) could be identified with no sequence homologies in the *E. coli* K-12 genome. These could be further sub-categorized in ‘uncharacterized proteins’ (53), ‘putative uncharacterized’ (5), ‘(DUF)’ domain containing proteins (21) and ‘UPF’ (HPs with unknown protein families, 3). 13 HPs had the characteristic ‘Protein Y…’ naming scheme. The distribution of those 95 HP-encoding genes within the genome can be found in the Supplementary (*Supplementary Figure 1*).

### iModulon generation and analysis

In order to investigate the functions for the 95 uncharacterized HP-encoding genes, publicly available RNAseq datasets from *E. coli* K-12 substrain MG1655 were downloaded from NCBI and quality filtered. 779 high-quality datasets from the MG1655 substrain were then used as input for the OptICA algorithm [27]. For the substrain BW25113, a separate OptICA analysis was not conducted due to the very small size of the dataset. The iModulons were identified by utilizing the PyModulon package, specifically employing the *Inferring iModulon Activities* function [32] for the BW25133 substrain. In total, 131 iModulons were obtained for the entire *E. coli* K-12 transcriptome. Three single gene iModulons were discarded from the analyses, since they were considered a result of artificial knockouts or overexpression of single genes in the dataset. Therefore, a total of 128 iModulons were selected for further analysis. Functional information on the HP-encoding genes was obtained by mapping the iModulons to existing experimentally derived regulons (groups of genes regulated by the same transcription factor), using data from the EcoCyc and RegulonDB databases. Additionally, Gene Ontology (GO)-based functional annotations for member genes of an iModulon (co-regulated gene sets) were identified using the PANTHER tool (Protein ANalysis THrough Evolutionary Relationships) [33]. A total of 51 HP-encoding genes were clustered into iModulons using OptICA. Of these, 46 genes were assigned putative functions based on information from regulator proteins and/or Gene Ontology (GO) categories. (**Figure 2**). Two genes, *yfeS* and *yffL* had information on their regulators (RpoE and NarL, respectively) but no assigned GO categories, while five genes (*ybaA, yfaP, yjgZ, ymgI and ymgJ*) (white) could not be associated with a regulator or a GO category. 44 of the HP-encoding genes could not be clustered into distinct iModulons (grey) (**Figure. 2**).

**Figure 2.**
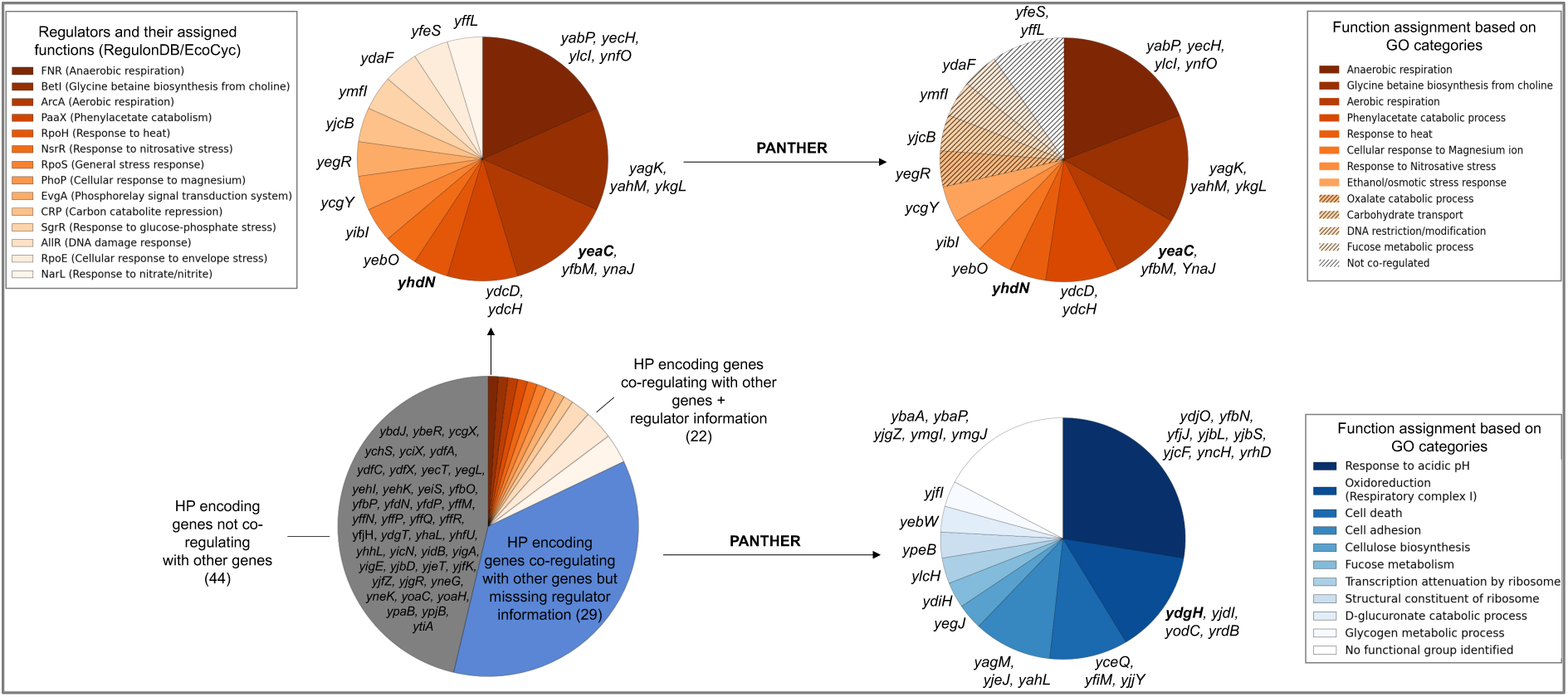
Regulator and functional classification for 95 HP-encoding genes in *Escherichia coli K-12*. 44 HP-encoding genes could not be clustered by the OptICA method [27] (grey), since they did not co-regulate with any other genes. 22 genes were characterized based on information on their regulators as obtained from RegulonDB and/or EcoCyc, as well as by Gene Ontology (GO) categories based on the co-regulating genes using the PANTHER tool (Protein ANalysis THrough Evolutionary Relationships) (orange) [33]. 29 genes co-regulated with other genes, but no information on their regulators could be obtained (blue). 24 out of these 29 genes could be assigned putative functions based on GO annotations of co-regulating genes derived from the PANTHER tool. The striped regions denote genes where the regulator-associated function from EcoCyc/RegulonDB did not match a GO-derived functional category obtained from PANTHER. Highlighted in bold are the three HP-encoding genes which were selected for *in vitro* testing.

In addition to the 95 HP-encoding genes, all other *E. coli K-12* genes were also analysed using this method. 2630 of the 2905 (91%) genes which have experimental evidence of function could also be characterized into iModulons. 1697 (64%) co-regulated with other genes, i.e. they were part of characterized iModulons with information on their regulator on RegulonDB and/or EcoCyc, while 933 of the genes were part of uncharacterized iModulons. For genes with known regulators, only 121 (5%) had either a non-matching functional category when compared with the PANTHER categories or a non-significant (*p*>0.05) functional category, and 95% had a matching PANTHER GO category. For genes which were part of uncharacterized iModulons, with no regulator information, 546 (58%) could be functionally classified using PANTHER. 1061 of the 1403 putative HP-encoding genes (75%), i.e. genes with functional characterisation based on sequence homology information, could be characterized into iModulons. 342 genes (25%) did not co-regulate with other genes. For genes that had co-regulation, 539 (51%) had information on their regulator on RegulonDB and/or EcoCyc. The other 522 (49%) were part of uncharacterized iModulons. For genes with known regulators, only 20 genes (4%) had either a non-matching function when compared with the PANTHER categories or a non-significant (*p*>0.05) functional category in PANTHER. For genes with no information on their regulator, 366 (67.9%), could however be characterized by PANTHER. Hence, about 95% of the iModulon functional categories matched the PANTHER GO categories, demonstrating the robustness of the method. Moreover, the inclusion of experimentally characterized genes within uncharacterized iModulons shows potential for the discovery of new regulons or co-regulated genes.

### Functional analysis of HP-encoding genes using *in silico* tools

All 95 HP-encoding genes were analysed based on multiple criteria including the generated iModulons from the OptICA algorithm, functional information based on the regulators of the co-regulated gene sets obtained *via* RegulonDB/EcoCyc and their GO categories from PANTHER. Information on metadata conditions pertaining to the highest level of gene expression relative to the reference gene *frr* (encoding the ribosome recycling factor) [42] and annotation of the most significant co-expressed gene were also extracted. Additionally, the sub-cellular localization, protein-protein interactions, genomic context, remote sequence homology as well as structural homology aided in forming hypothesises for assigning functions to the respective HPs. 33 of the protein encoding genes (highlighted in green) could be assigned functions with a high confidence. 28 of the genes were assigned functions based on lower confidence, since only information from a maximum of two sources could be well-correlated (highlighted in yellow). A clear inference could not be made for 34 of the genes (highlighted in red). Information retrieved for each gene and functional inference (if applicable) is detailed in **Table 1**. A pictorial overview of Table 1 based on the functional annotations for the 95 HPs can be found in *Supplementary Figure 2*.

**Table 1.**
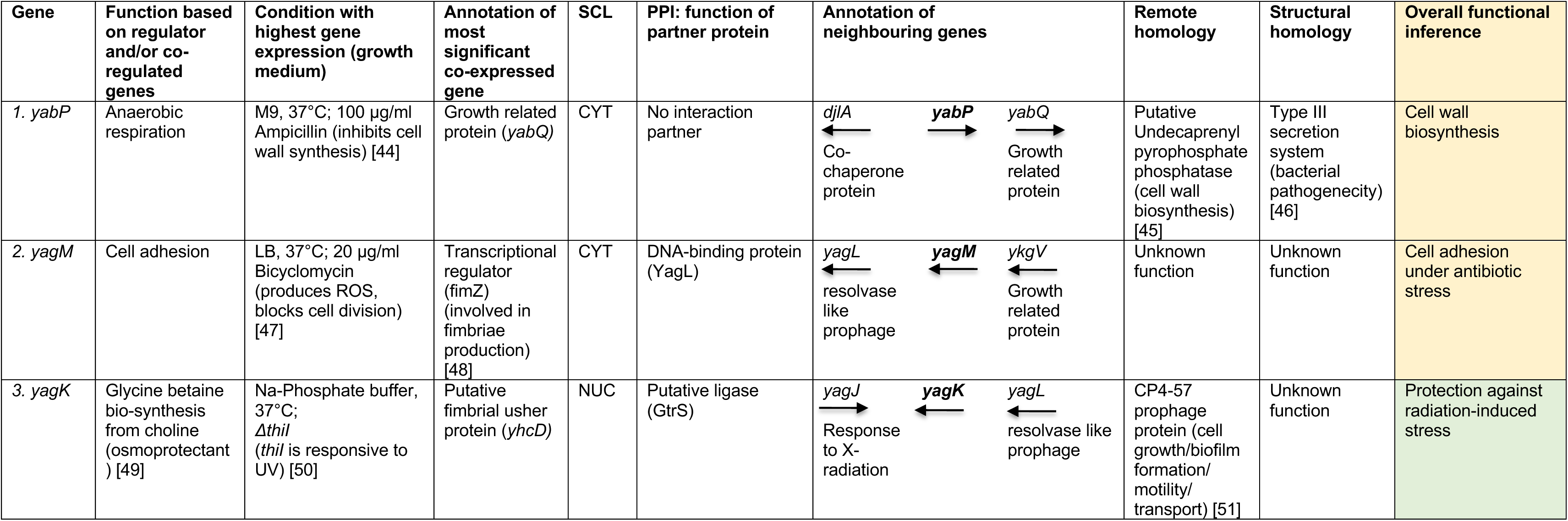
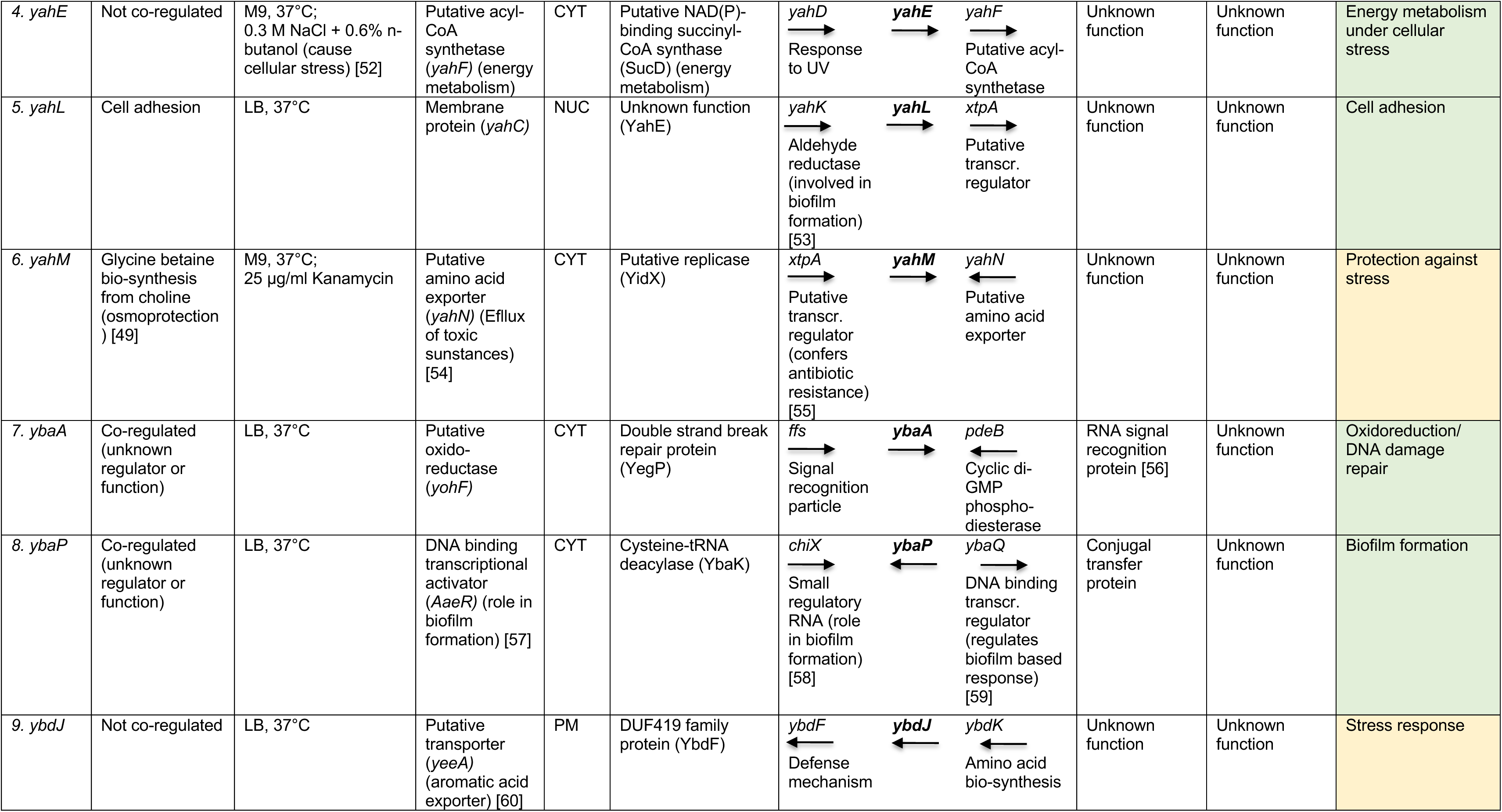
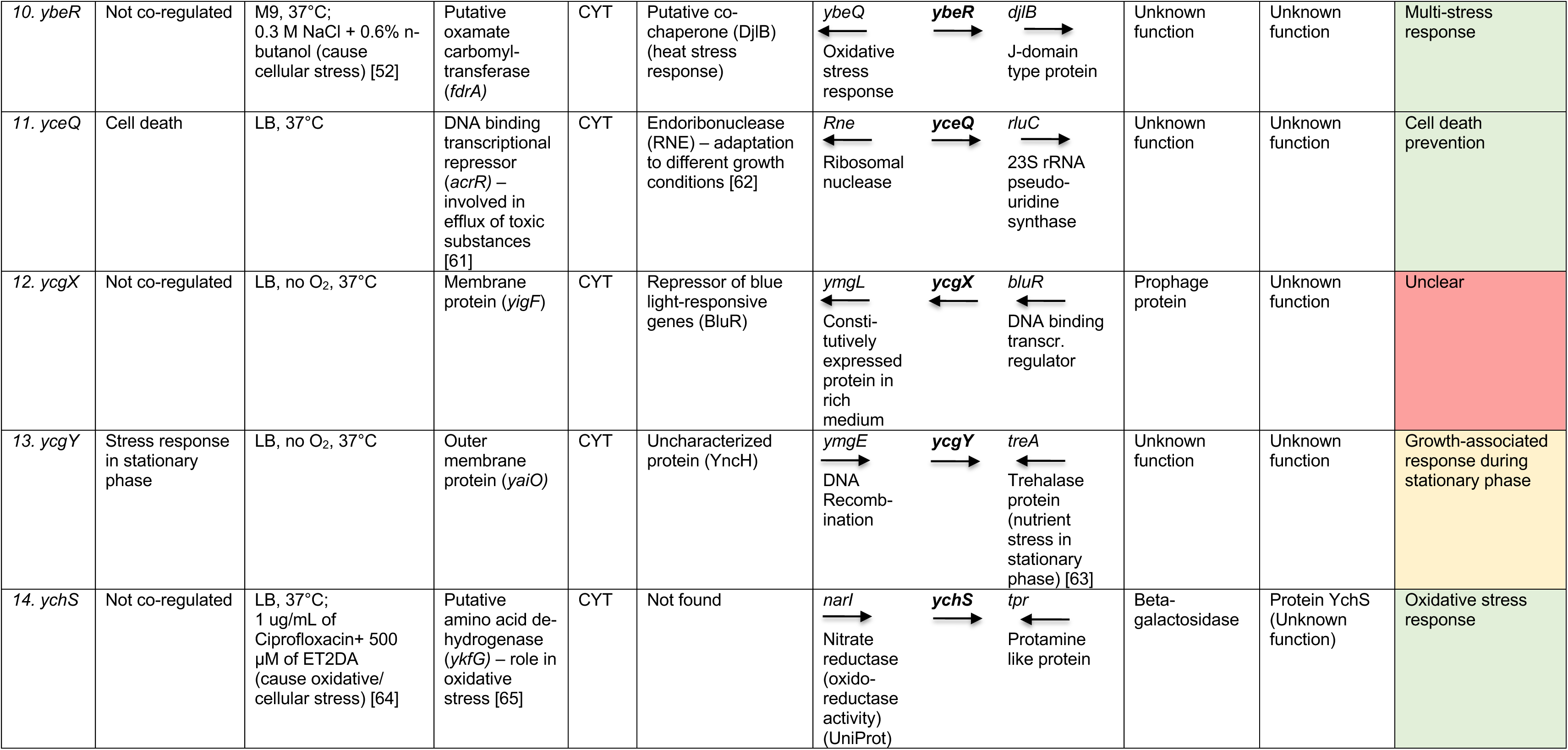
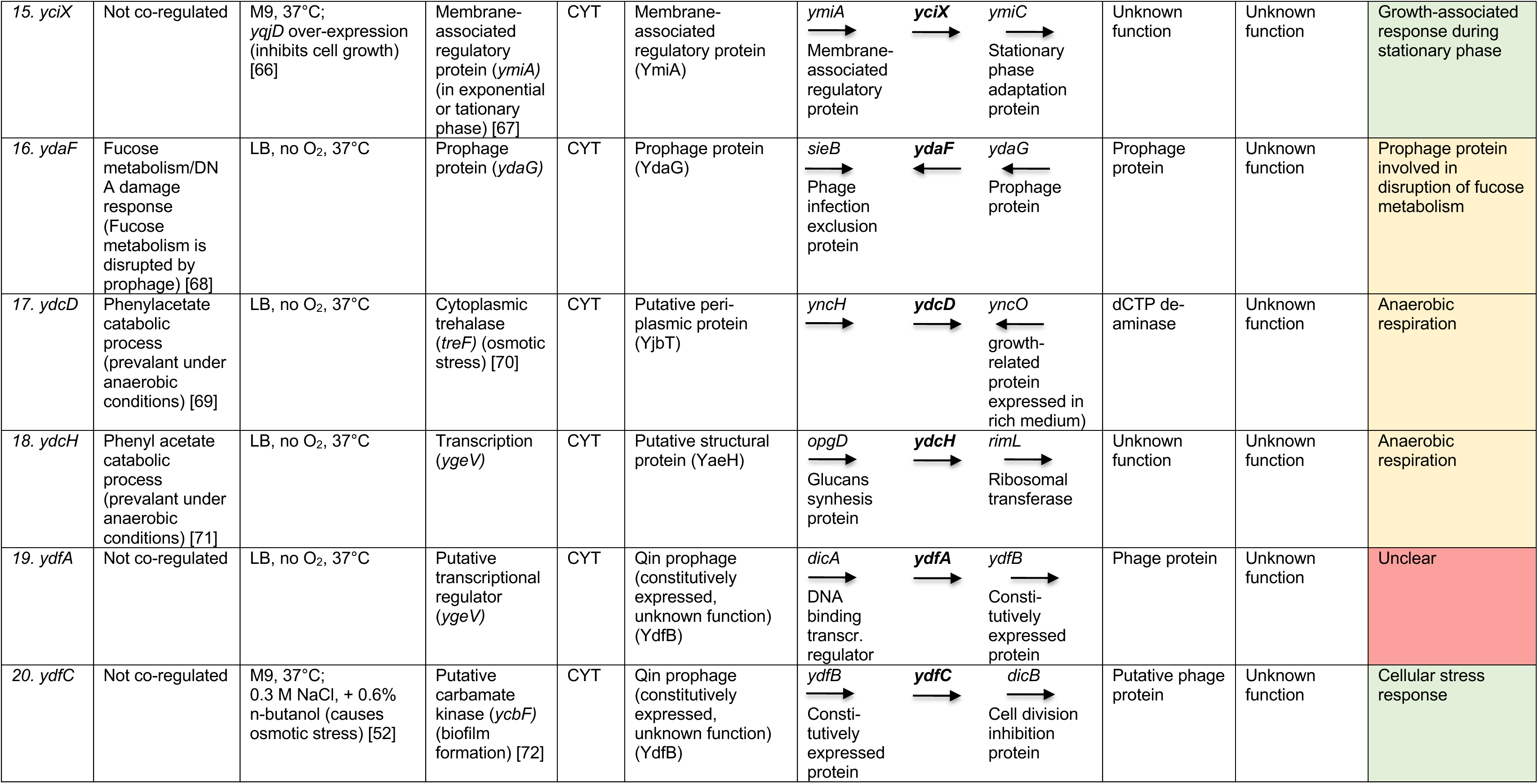
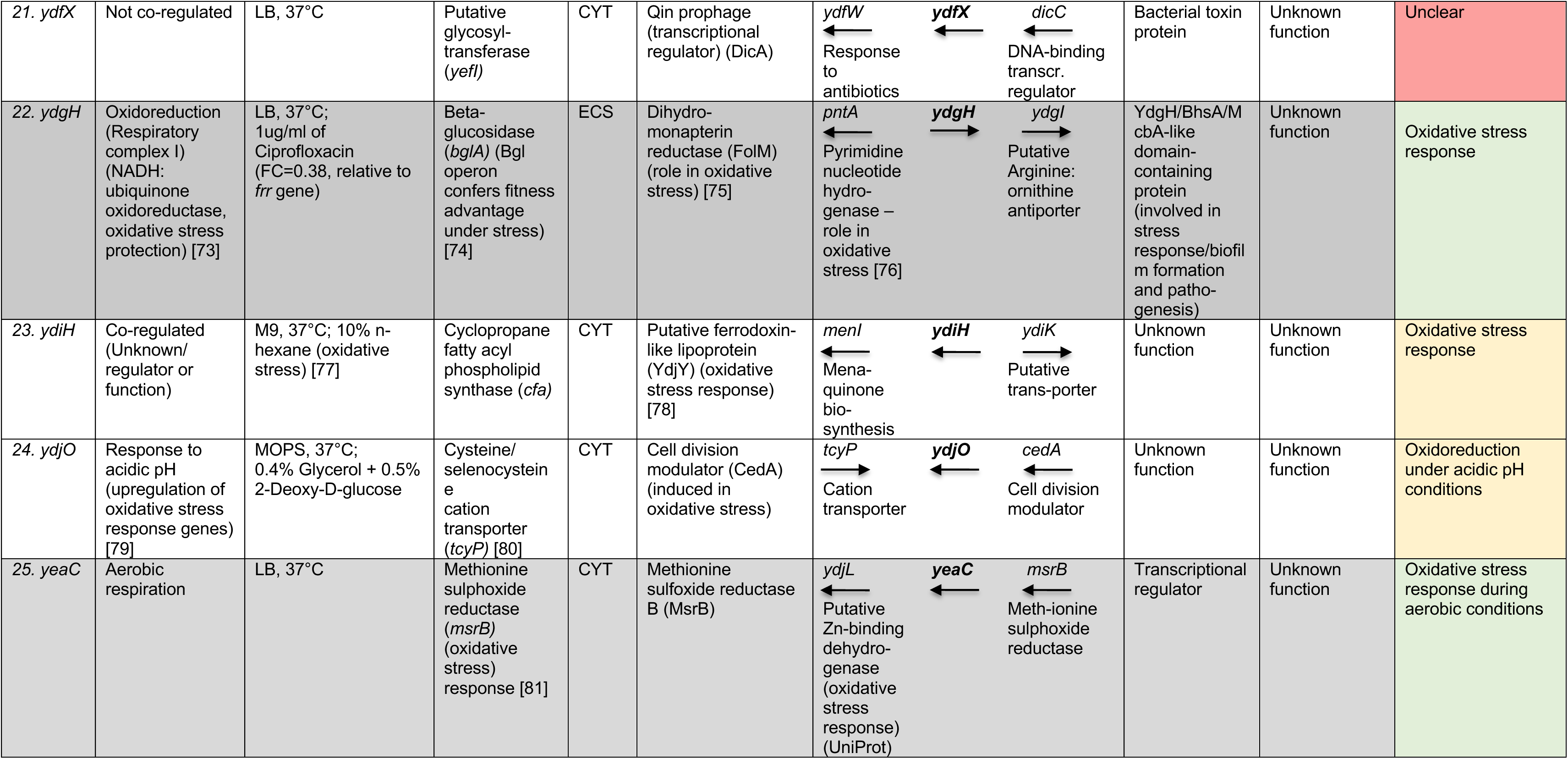
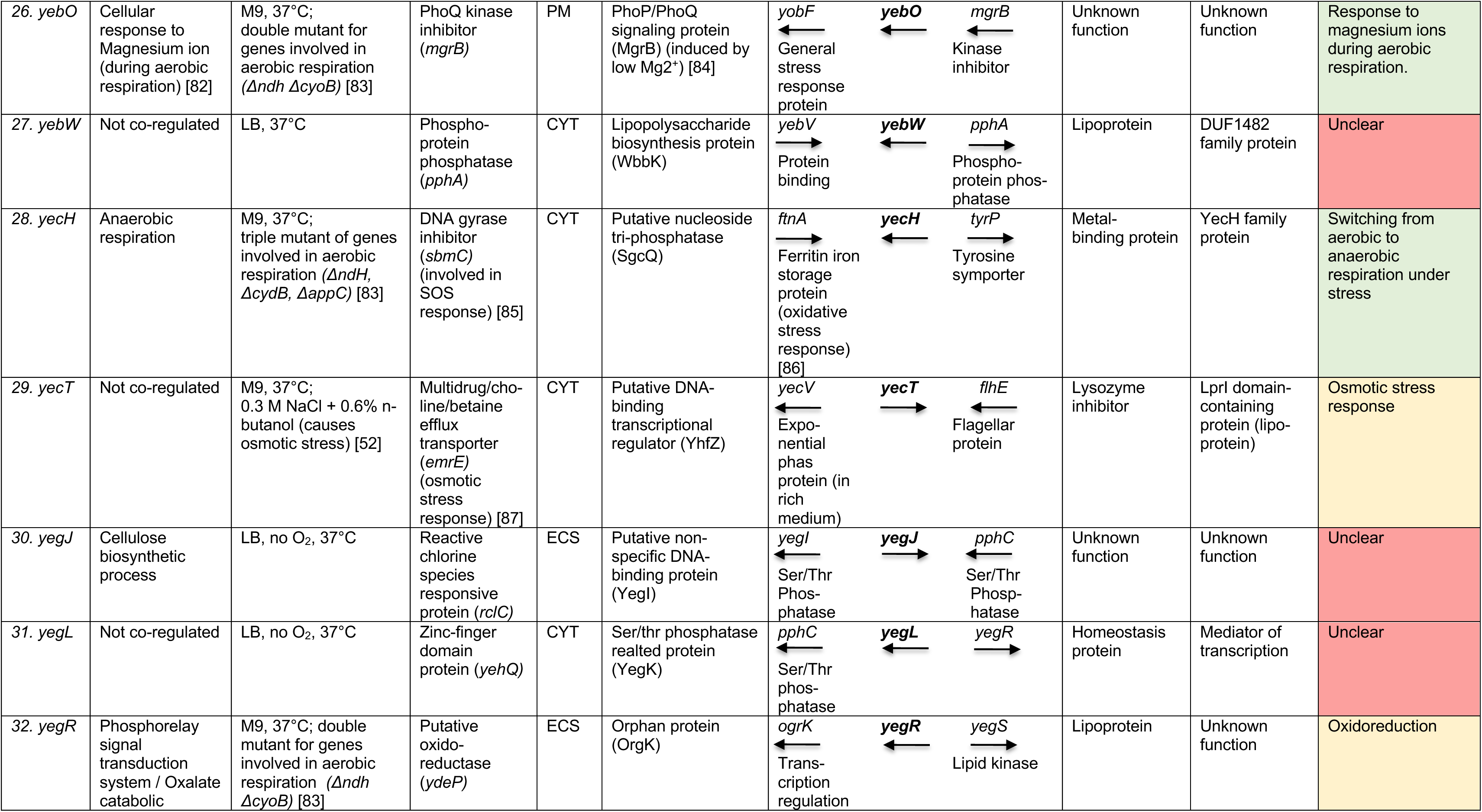
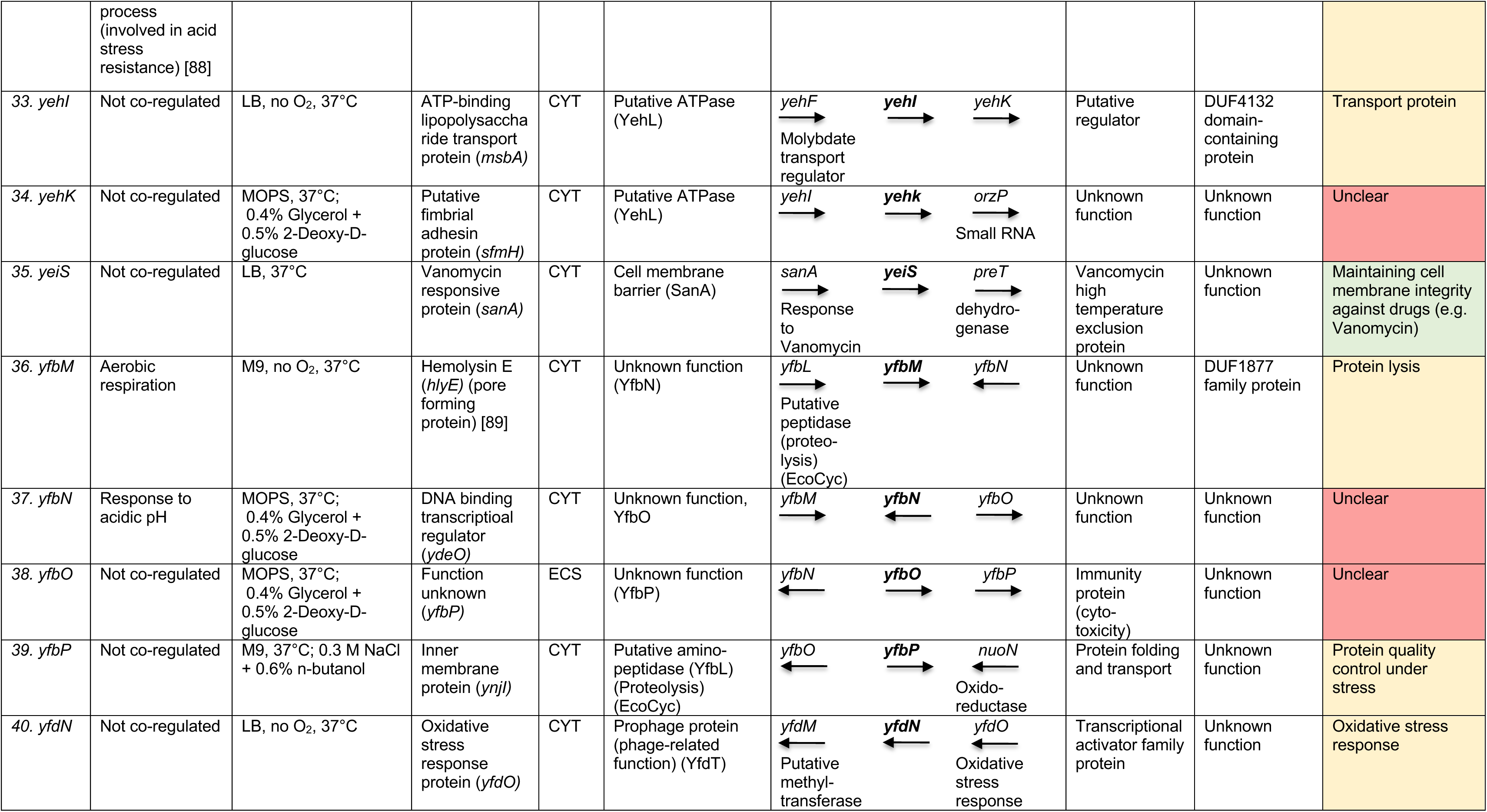
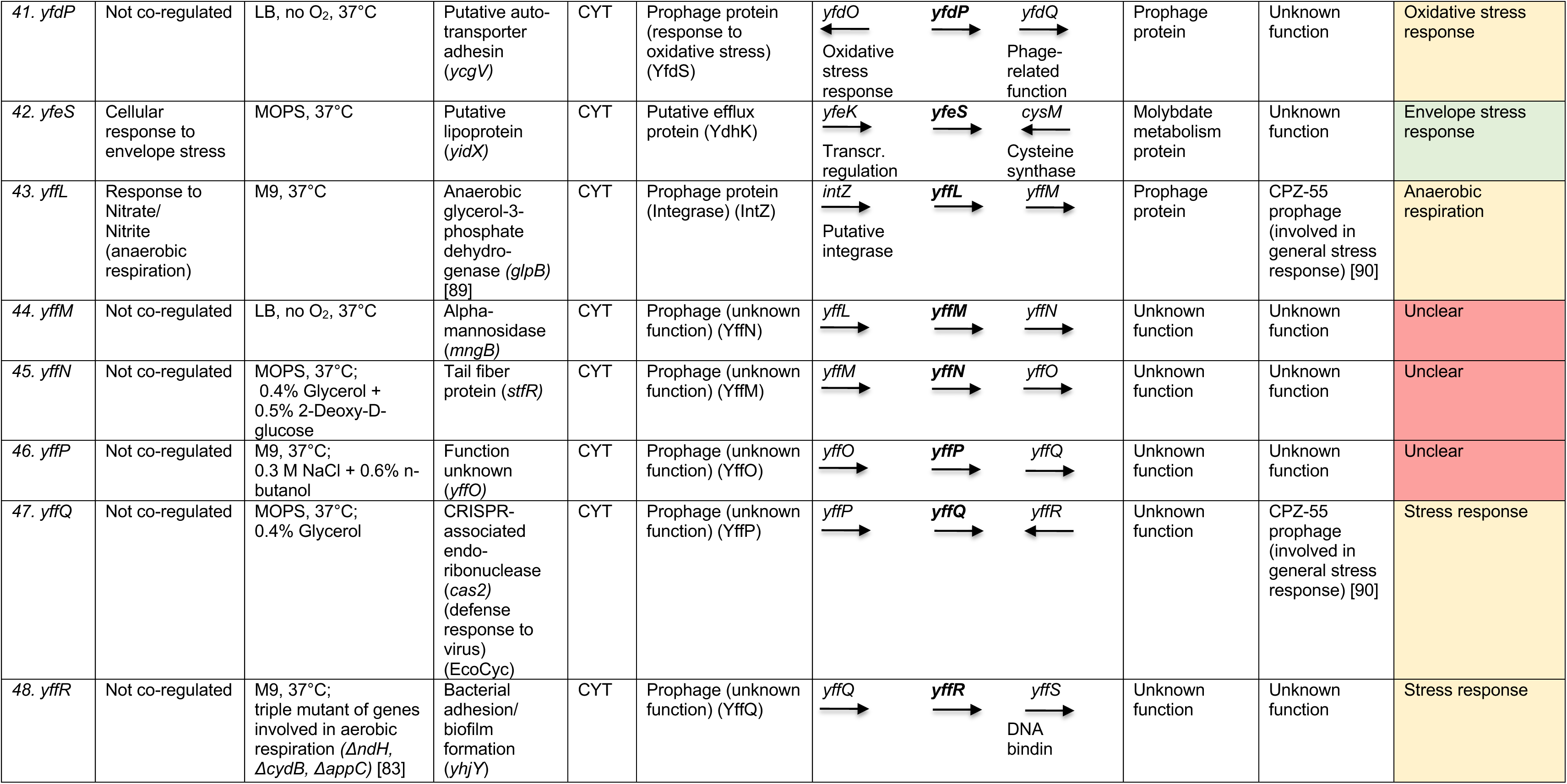
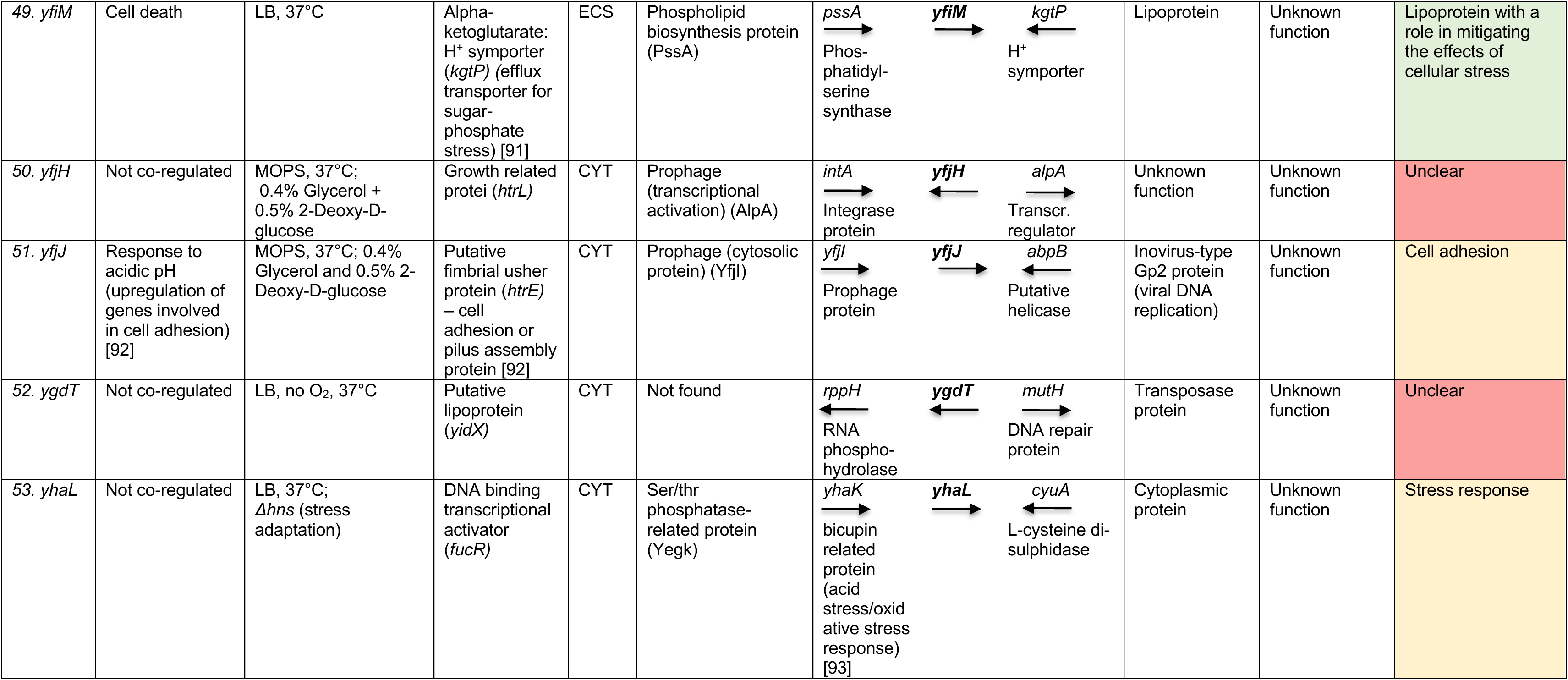
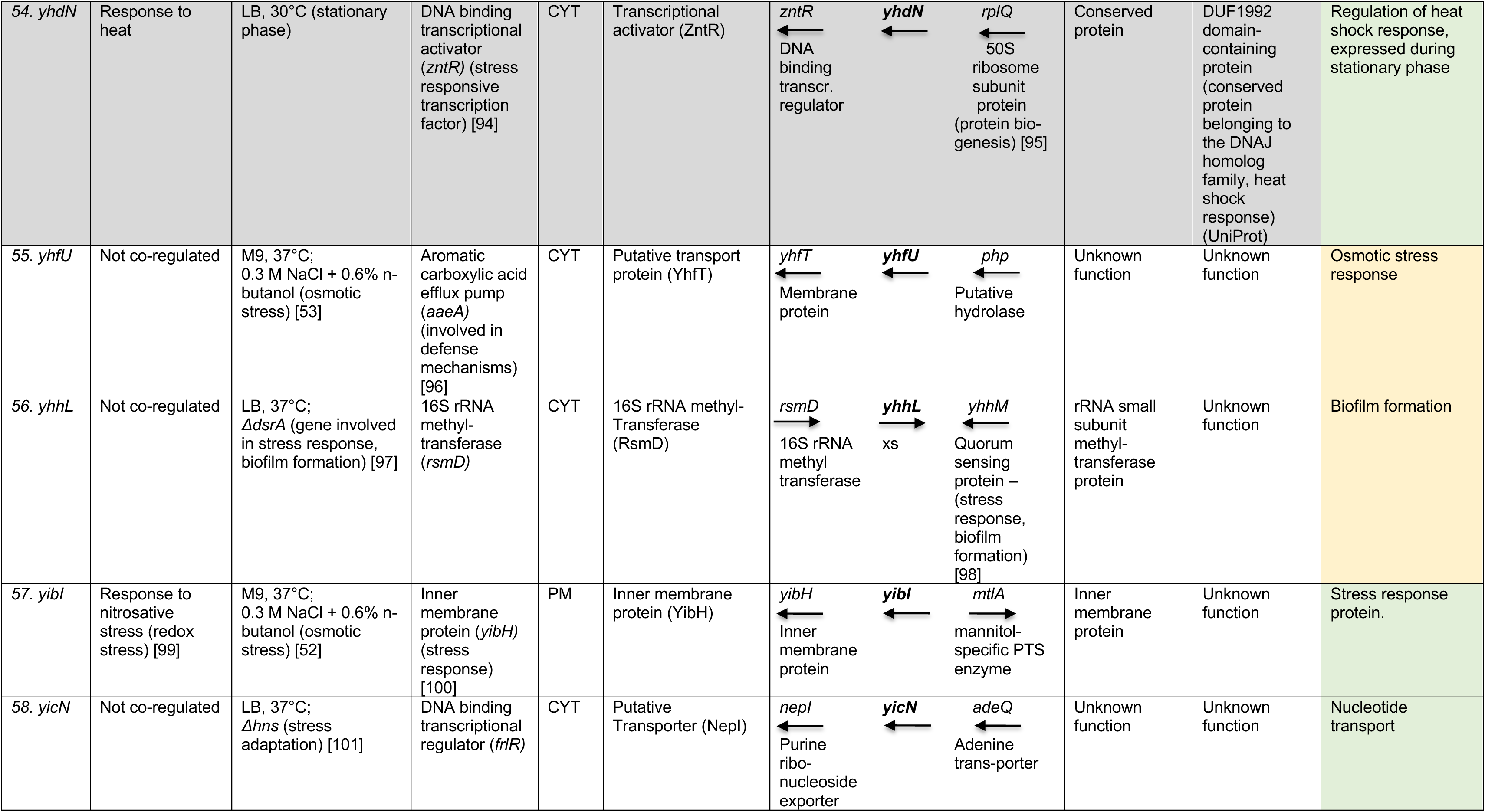
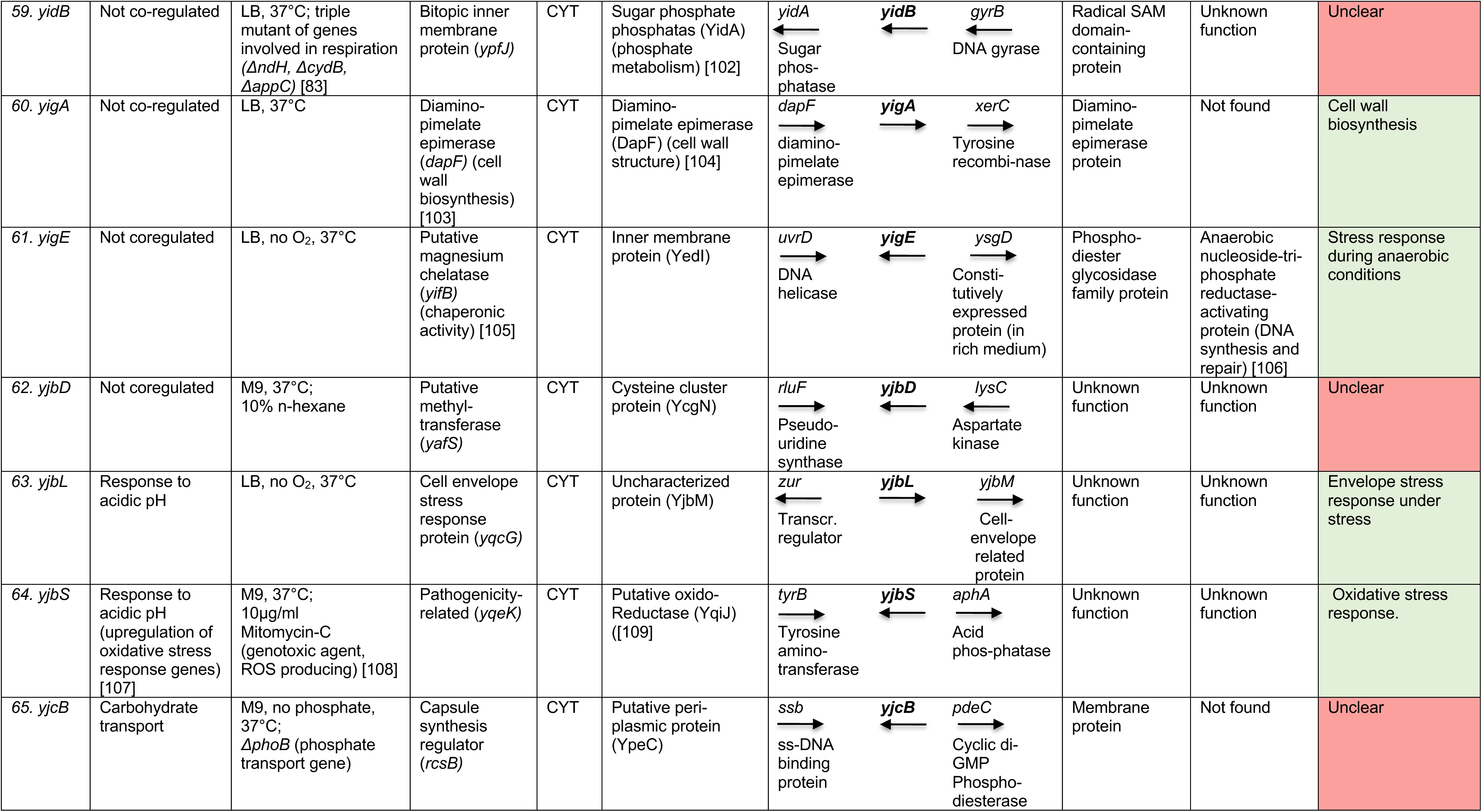
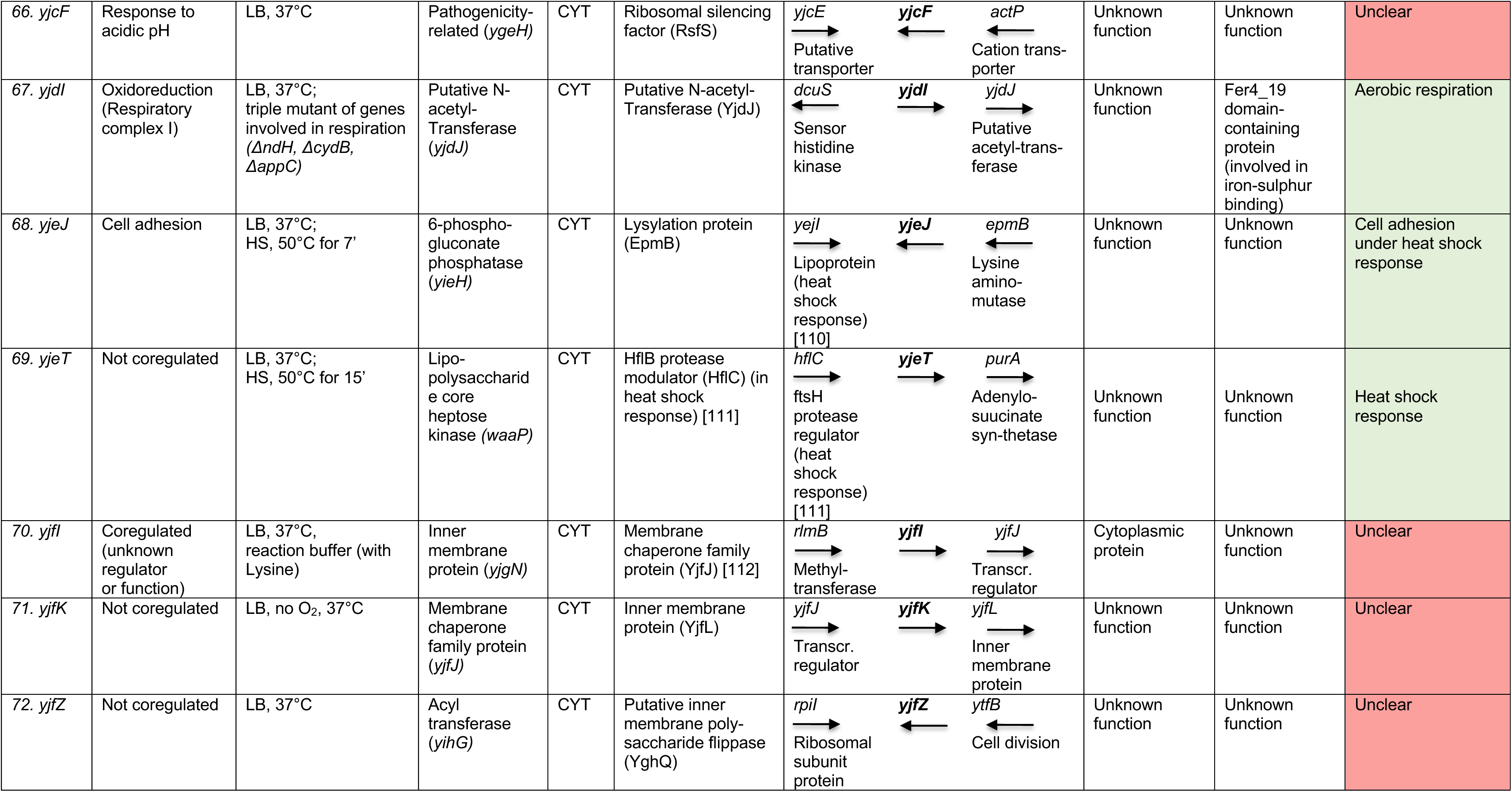
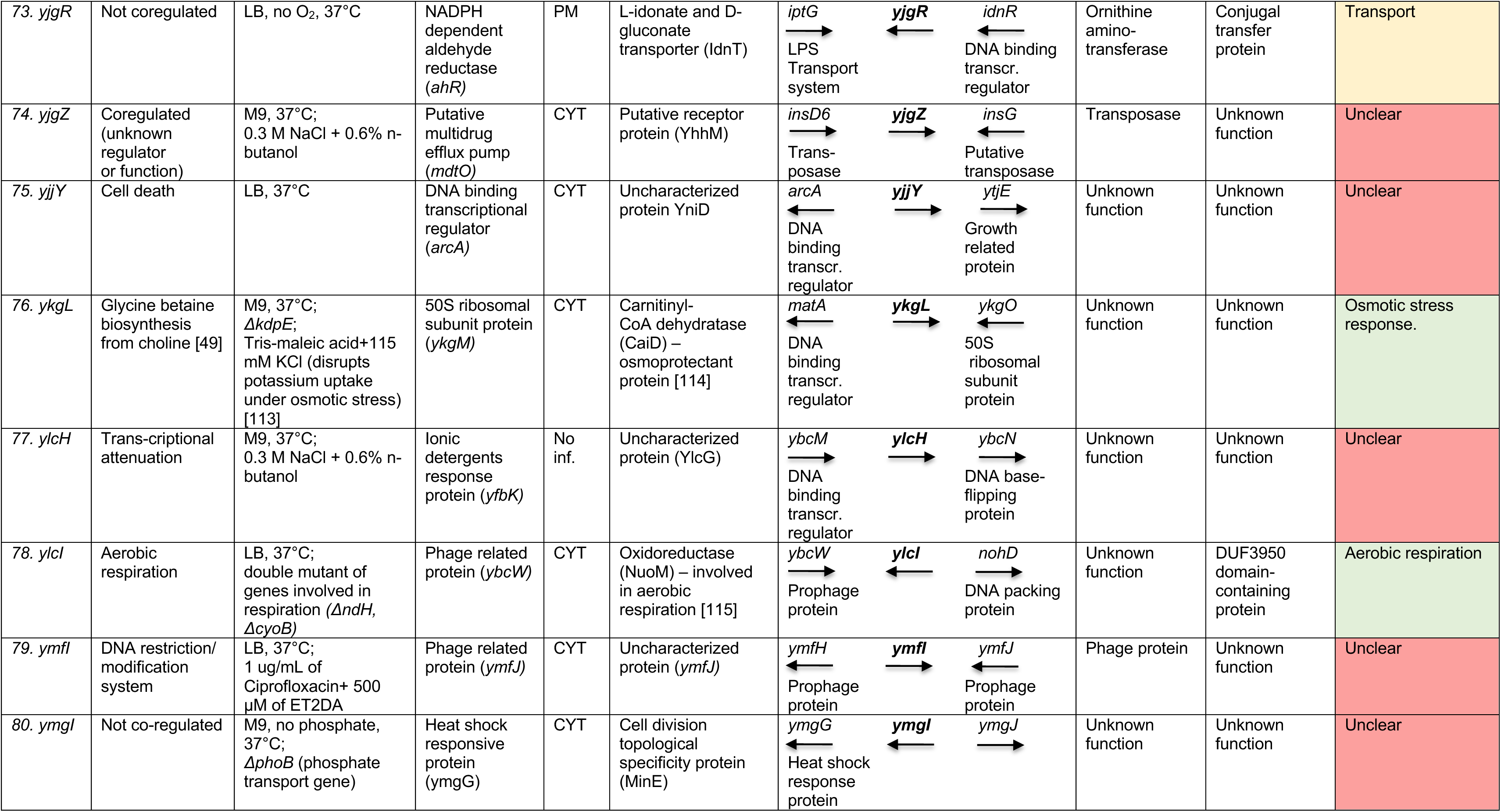
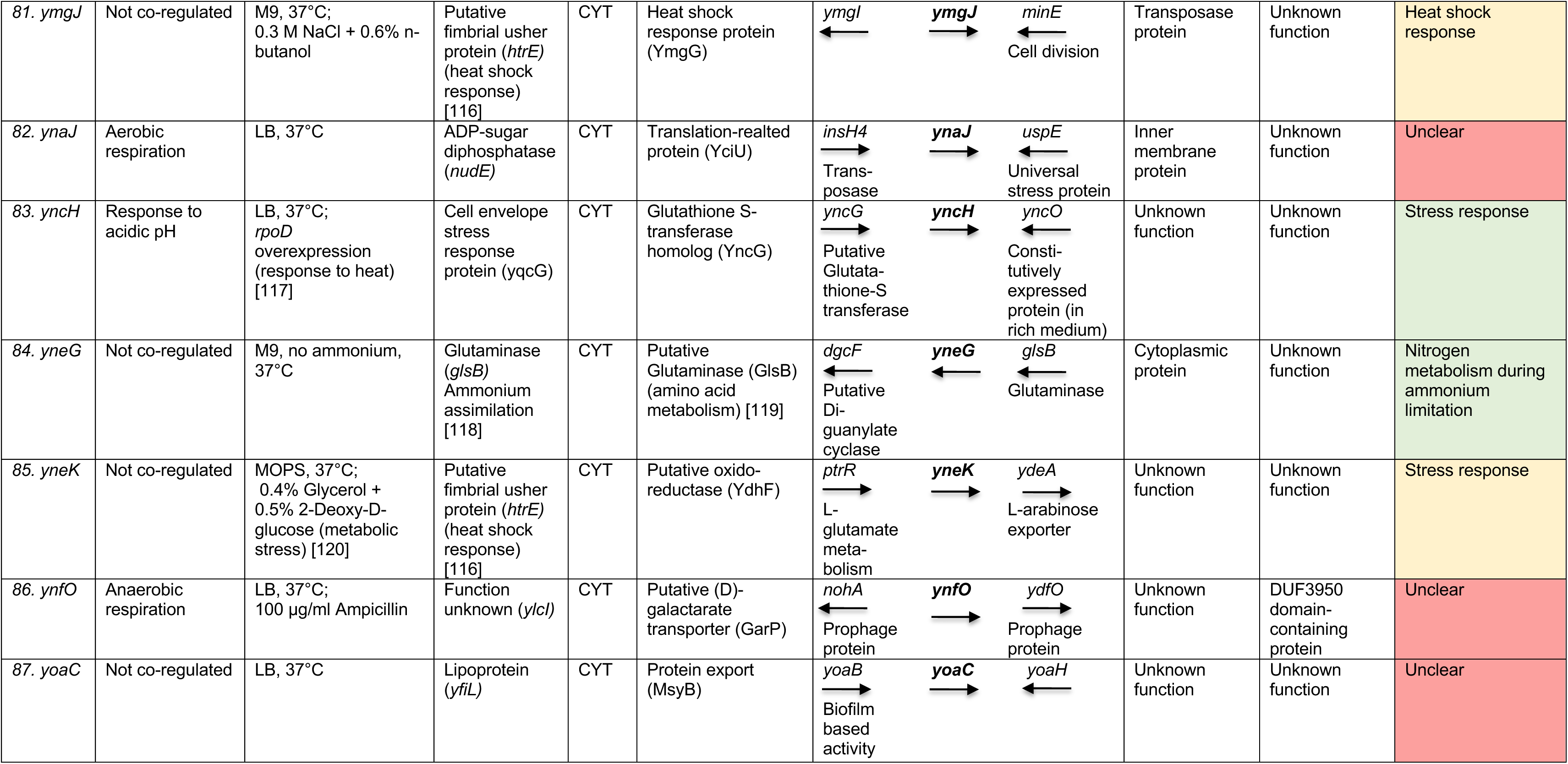
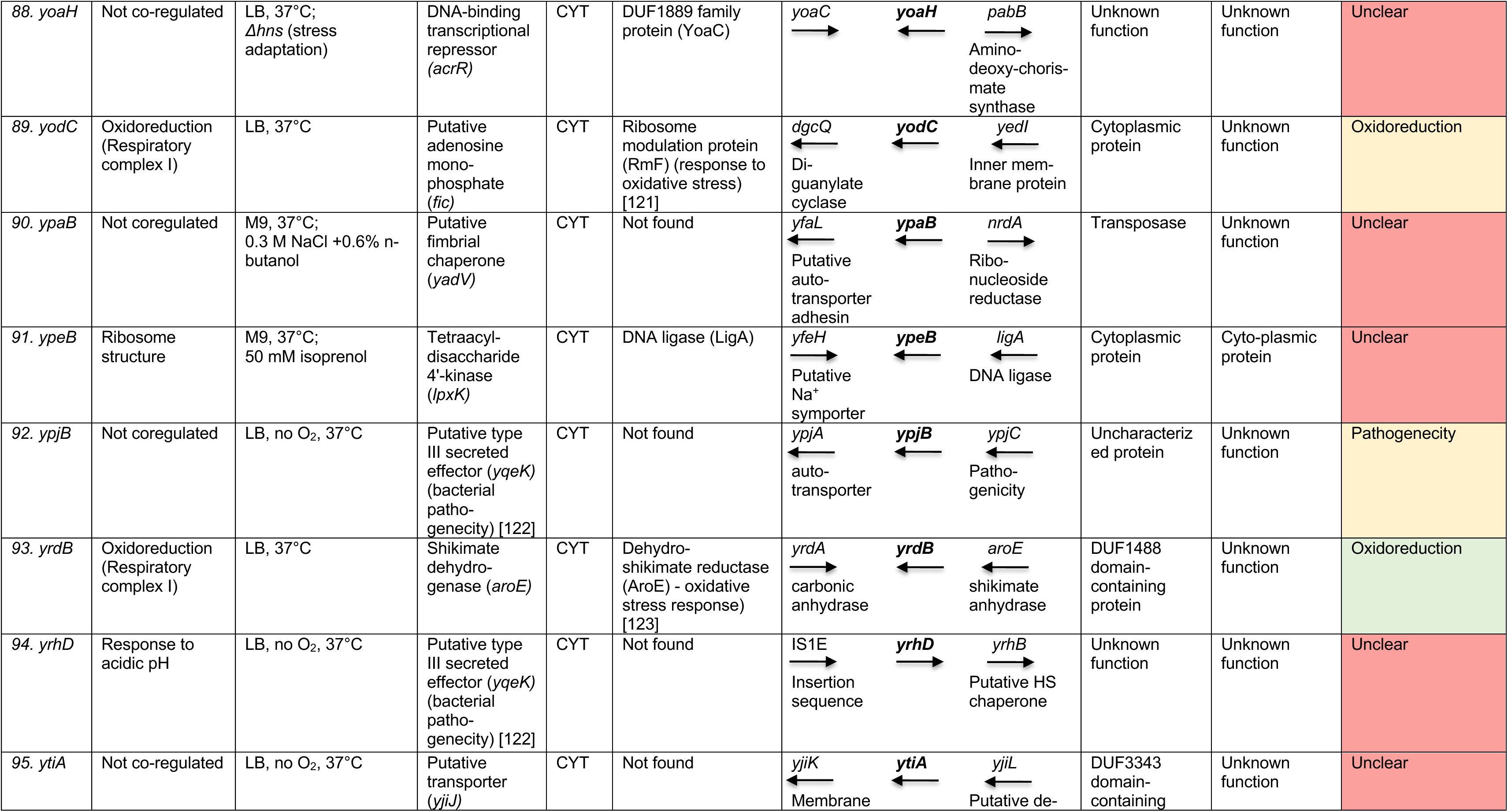
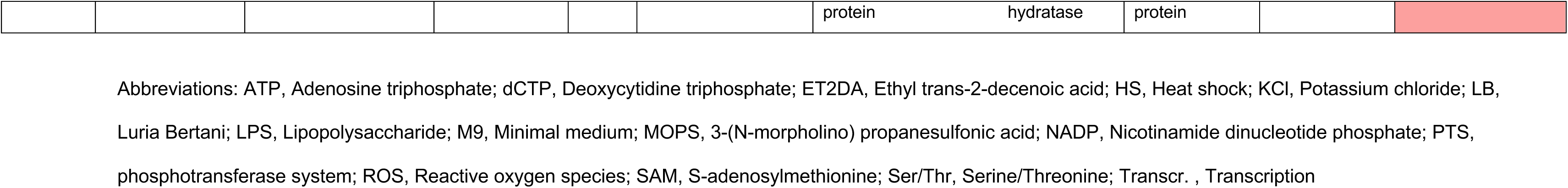
A comprehensive functional interference of HPs based on different *in silico* tools. Annotations are based on regulator information from RegulonDB/EcoCyc and/or GO category based on co-regulated genes obtained from PANTHER) [33](as shown in Fig. 2). The highest gene expression condition, obtained from the Tjaden dataset [43], with metadata conditions categorized by the type of medium, oxygen availability (normal O_2_ conditions unless stated otherwise), temperature etc. in comparison to the reference gene ‘*frr’* are described. Functional annotations of co-expressed genes were retrieved from EcoCyc [36]. ‘SCL’ or sub-cellular localization inferred from the BUSCA tool [34] are abbreviated as either ‘NUC’ for nuclear ‘CYT’ for cytoplasmic, ‘ECS’ for extra cellular space or ‘PM’ for plasma membrane. Functional information from protein-protein interaction (PPI) analyses was obtained from STRINGdb [35]. Annotations for neighbouring upstream (←) and downstream (→) genes in context of the HP-encoding gene were retrieved from the EcoCyc database. Additionally, remote homology was assessed using the Uniref30 database using the Hhblits tool [37], and structural homology was inferred from the AlphaFold Protein Structure Database [38]. A functional inference can be made for genes supported by multiple lines of evidence when similar functional information was provided by at least two sources. Proteins with well correlated information from at least three sources are highlighted in green (with higher confidence) and proteins with information that can be correlated using at least two sources are highlighted in yellow (with lower confidence). Protein encoding genes highlighted in grey were selected for *in vitro* testing. An unclear inference suggests that, based on the different knowledge bases, the extracted information did not provide enough confidence or could not be well correlated to assign a function (red).

Genes with a higher confidence in the inference section (green) were analysed for candidates to be used for *in vitro* testing. Three HP-encoding genes, namely *yhdN*, *yeaC*, and *ydgH*, were selected for further analysis (highlighted in grey) (**Table 1).**

### Protein encoding gene *yhdN*

Protein YhdN, characterized by the presence of a domain of unknown function (DUF1992), was predicted to be involved in heat shock response. The gene encoding *yhdN* is regulated by the RNA polymerase sigma factor RpoH (σ32), which serves as the primary heat shock transcriptional regulator. Under normal conditions, RpoH levels are kept low due to chaperone-mediated degradation; however, these levels increase significantly upon heat shock [124]. Gene Ontology (GO) annotation for this cluster is categorized under ‘response to heat’. Based on publicly available transcriptomic data [43], the *yhdN* gene exhibited significant expression in cells grown to the stationary phase, with a Log_2_FC of 5.5 relative to the reference gene *frr*, signifying a 45 times higher expression of *yhdN* gene. This observation suggests that the *yhdN* gene is highly expressed during the stationary phase and therefore might have a role in nutrient stress response. Additionally, *yhdN* expression was upregulated following exposure to a transient heat shock at 50°C for 15 minutes, with a Log_2_FC of 4.8 and 27 times higher expression, suggesting that the gene might also play a role in heat shock response. Co-expression analysis using Spearman correlation of the gene resulted in a high correlation with the gene *zntR* (0.93) encoding an HTH (helix-turn-helix)-type transcriptional regulator [125] which is involved in protein stability and prevention of aggregates [126,127]. Genes *yhdN* and *zntR* are also part of the same transcription unit (EcoCyc database). Other genes with high correlations to *yhdN* were *relB* (0.84)*, pspB* (0.84) *and pspC* (0.84), all of which are involved in stress response [128] (*Supplementary Figure 3A*.). Gene *relB* encodes for an antitoxin protein involved in regulation of growth [129] while genes *pspB* and *pspC* encode for phage shock proteins that are known to be induced by infection with bacteriophages or ethanol, osmotic and/or nutrient stress as well as heat shock conditions [130]. Protein-protein interaction (PPI) analysis showed highest interaction confidence with ZntR. The local genomic context analysis showed genes *zntR* and *rplQ* (a ribosomal subunit protein) as neighbouring genes. Protein YhdN showed remote sequence homology to a protein cluster comprising of uncharacterized conserved proteins and was structurally homologous to conserved proteins with similarity to J-domains of chaperones, regulating the activity of heat-shock proteins [131]. To elucidate the potential function of *yhdN*, we hypothesized that its absence would affect bacterial growth under heat shock conditions and possibly the cells stationary phase under nutrient limitation. This hypothesis was tested by comparing the growth patterns of the *E. coli* K-12 substrain BW25113 wild type and its isogenic *ΔyhdN* knockout mutant under a transient heat shock condition (50°C) and recovery at 37°C in LB and nutrient-limited M9 medium.

### Protein encoding gene *yeaC*

Protein YeaC is characterized by the presence of a DUF1315 domain. Gene *yeaC* is predicted to be governed by the regulator ArcA, which regulates a group of proteins involved in redox homeostasis and aerobic respiration [132]. The Gene Ontology (GO) annotation for this cluster was predicted as ‘aerobic respiration’. Analysis of gene *yeaC* in publicly available transcriptomic datasets showed slightly higher expression levels against gene *frr* in LB medium at 37°C under normal O_2_ conditions (log_2_ FC 1.9) (**Table1**) and in knock-out mutants for genes encoding type II NADH:quinone oxidoreductase (*ndh*), Cytochrome bd-I ubiquinol oxidase (*cydB*) and Cytochrome bd-II ubiquinol oxidase (*appC*), involved in the Tricarboxylic acid cycle or oxidative phosphorylation [133] (log_2_ FC 1.7). Based on the co-expression analysis of public data, the highest correlation was observed with a gene encoding for a peptide methionine sulfoxide reductase (*msrB),* which is an oxidoreductase and known to be highly expressed during oxidative stress to maintain the cell-redox homeostasis [134] (*Supplementary Figure3B.*). Other highly correlated genes involved succinate dehydrogenase cytochrome b556 subunit (*sdhC)*, succinate dehydrogenase hydrophobic membrane anchor subunit (*sdhD)* and Fumarate hydratase class I, aerobic (*fumA),* which are all involved in the tricarboxylic acid or TCA cycle [135,136] (*Supplementary Figure 3B*.). PPI analysis also showed highest interaction confidence with MsrB. Additionally, it neighbours the *msrB* gene and a putative zinc-binding dehydrogenase, *ydjL* which is also involved in oxidoreduction (UniProt) (**Table 1**). Protein YeaC shows remote sequence homology to other proteins involved in transcriptional regulation. Structural homology revealed similarities to a group of uncharacterized proteins with unknown functions. We hypothesized that *yeaC* affects the cell-redox homeostasis and cellular growth under oxidative stress. This hypothesis was tested by comparing the growth patterns of the *E. coli* K-12 substrain BW25113 wild type and its isogenic *ΔyeaC* knockout mutant with a sub-lethal concentration of H_2_O_2_ (2.5 mM) [137].

### Protein encoding gene *ydgH*

Protein YdgH has a DUF1471 domain of unknown function. It is predicted to be part of an unknown cluster, with no information on its regulator. The GO annotation for this cluster, however, is categorized under oxidoreduction (complex I). The energy-converting NADH:ubiquinone oxidoreductase respiratory complex I, is the main entry point for electrons from NADH into the respiratory chains in bacteria [138]. Based on the public datasets, we could not obtain a gene-specific condition where gene *ydgH* had higher expression compared to gene *frr.* Co-expression analysis of *ydgH* from the publicly available RNAseq datasets revealed highest correlation with a 6-phospho-beta-glucosidase (*bglA*) encoding hydrolase, involved in the hydrolysis of phosphorylated beta-glucosides [139]. Other highly correlated genes were phosphoglycolate phosphatase *(gph)* [140] and ribulose-phosphate 3-epimerase *(rpe)* [141], involved in oxidative stress response (*Supplementary Figure 3C*.). PPI analysis showed highest interaction confidence with YjfY, a putative HP, with a possible role in stress resistance (UniProt). Genomic context provided information on a neighbouring upstream gene involved in NAD(P)^+^ transhydrogenase activity, annotated as NAD(P) transhydrogenase, *pntA*, which is known to play a role in oxidative stress [142]. The protein showed remote sequence homology to a YdgH/BhsA/McbA-like domain involved in stress response/biofilm formation and pathogenesis [143]. The structural similarity was only to uncharacterized proteins with unknown function. Following a similar hypothesis to that proposed for *yeaC*, we evaluated the growth patterns of the wild type and its isogenic *ΔydgH* knockout mutant under oxidative stress.

### *In vitro* analyses of candidate proteins

#### Effect of heat shock on growth

To determine the effect of a transient heat shock at 50°C (7 minutes), the wild type and the mutant *ΔyhdN E. coli* BW25113 strains were grown in LB medium. The *yhdN* mutant was seen to have a growth defect of ∼18% after 6h of growth based on the OD values obtained (**Figure 3A**). It was observed that after 6h of growth, the mutant could not resume normal growth compared to the wild type. Additionally, the effects of a transient heat shock in Nitrogen-limited M9 medium showed a significant growth defect of ∼50% (**Figure 3B**). Growth curves for WT and *ΔyhdN* in different temperatures can be found in the supplementary (*Supplementary Figure 4A*.). Growth defects were also observed in the *yhdN* mutant compared to the wild type when both the strains were grown continuously at 50°C. Since growth at 50°C is however severely inhibited and cell started dying after 3h (*Supplementary Figure 4B*.), it was difficult to say whether this was solely based on the absence of the protein YhdN in the mutant cells.

**Figure 3.**
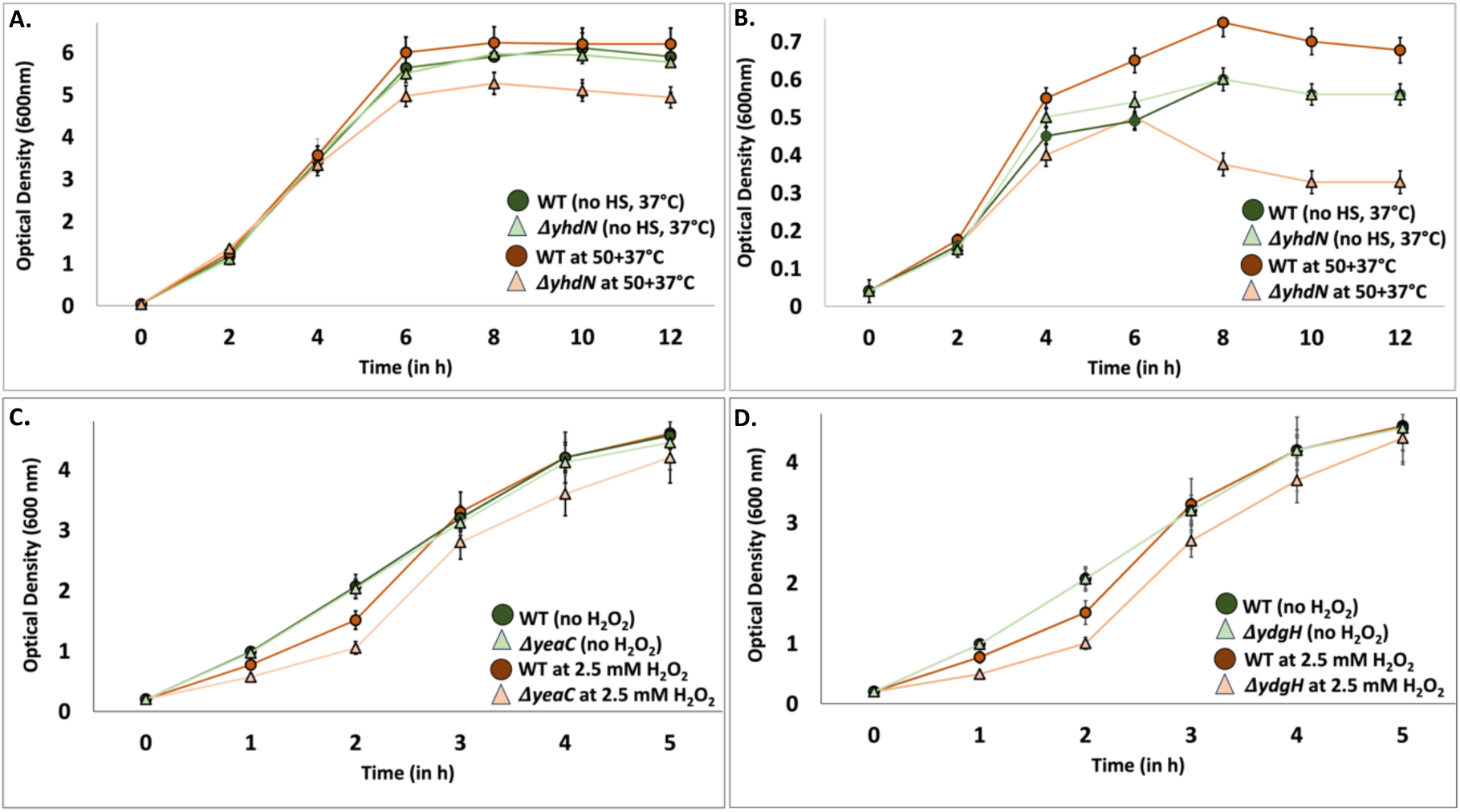
Growth curves of *E.* coli K-12 BW25113 wild type (WT) and isogenic deletion mutants. (A, **B)** Effect of transient heat shock on WT and **Δ*yhdN* cells. WT and the respective mutant strain were exposed to a transient heat shock at 50°C for 7 minutes, (at OD600 nm = 0.04) and grown for 12 h in **A.** LB medium and **B.** in Nitrogen-limited M9 medium. **(C, D)** Bacterial growth of WT and respective mutants at 37°C exposed to sub-lethal concentration of 2.5 mM H2O2, added during the exponential phase (OD600 = 0.2) and grown for 5h. **C.** WT and **Δ*yeaC* and D. WT and **Δ*ydgH* cells. WT are indicated by filled circles (●) and mutants by filled triangles (▲). Green lines represent controls, orange lines stress conditions. Average of three independent readings taken for each specified condition. Error bars on the graph indicate standard deviation from the mean.

#### Effect of sublethal H_2_O_2_ concentration on growth

To determine the effect of sublethal H_2_O_2_ concentration, the wild type (WT), *ΔydgH* and *ΔyeaC E. coli* BW25113 strains were grown in LB medium until an OD of 0.2 and then exposed to an H_2_O_2_ concentration of 2.5 mM, which is known to causes DNA damage and oxidative stress [143] (**Figure 3C & D**). To study the responses due to oxidative stress, the absorbance values (OD_600_) after each hour over a 5h time period were measured. H_2_O_2_ was added to the cell culture during the exponential phase of growth, which is characterized by rapid cell division and metabolic activity and where the quorum-sensing signalling, i.e. process which allows the bacteria to communicate and adjust gene expression according to cell density, is minimal. Under non-stress conditions, the WT and mutant strains (*ΔydgH* and *ΔyeaC*) exhibited comparable growth, with no significant differences observed. A growth lag was observed in the mutants (*ΔydgH* and *ΔyeaC*) as compared to wild type when the medium was supplemented with 2.5 mM H_2_O_2_. A growth defect of ∼25% was observed in the mutant when WT was compared to *ΔyeaC* at the 1h timepoint. This was reduced to ∼7% after 5h of growth underscoring the probability of the gene being expressed at this time point to counterfeit the oxidative stress. Similarly, a growth defect of ∼36% was observed in *ΔydgH* as compared to WT at 1h, which decreased to ∼2% after 5h, again probably due to stress adaptation and expression of gene *ydgH*.

### Differential gene expression analysis

For a comparative analysis of differentially expressed genes (DEGs) and their functions in WT *vs.* the mutant strains, all up- and down-regulated genes were analysed (*Figshare > DEGs > Table1*). For positively regulated DEGs, representing genes upregulated in the WT strain, a Log_2_ fold change (Log_2_FC) threshold of ≥1.5 was applied, corresponding to a minimum 2.8-fold increase in expression levels. This cut-off ensured the inclusion of genes with significant upregulation while minimizing noise [144]. Conversely, for negatively regulated DEGs, corresponding to genes upregulated in the mutant strains, a Log2FC threshold of ≤-1.5 was used.

#### WT *vs. *Δ*yhdN*

Analysis of the WT strain at the 2h and 6h time points showed that *yhdN* was only significantly expressed after 6h, hence this time point was chosen. In total, 484 DEGs (*p_adj_*≤0.1) were identified, (181) 213 genes with a positive Log_2_ FC and 271 with a negative Log_2_FC of 1.5. Within the positive DEGs, an analysis of the top 10 genes, resulted in one gene involved in protein folding, five in stress response and four in motility/chemotaxis (**Figure 4A**). Significantly upregulated genes in the WT strain were *tdcB* (Log_2_ FC=7.37) and *pyrB* (Log_2_ FC=6.14). *tdcB*, encoding for threonine dehydratase is known for being expressed during nutrient deprivation (EcoCyc) and gene *pyrB* is involved in pyrimidine metabolism and associated with motility under stress [145]. Gene *tnaA* (Log_2_ FC=6.02), which encodes for tryptophanase, has been recently studied for its role in protein folding and dissolution of protein aggregates [146]. Motility and chemotaxis genes such as *fliL* (Log_2_ FC=6.24)*, flgD* (Log_2_ FC=5.99) are also known to be involved in response to environmental stresses [147].

**Figure 4.**
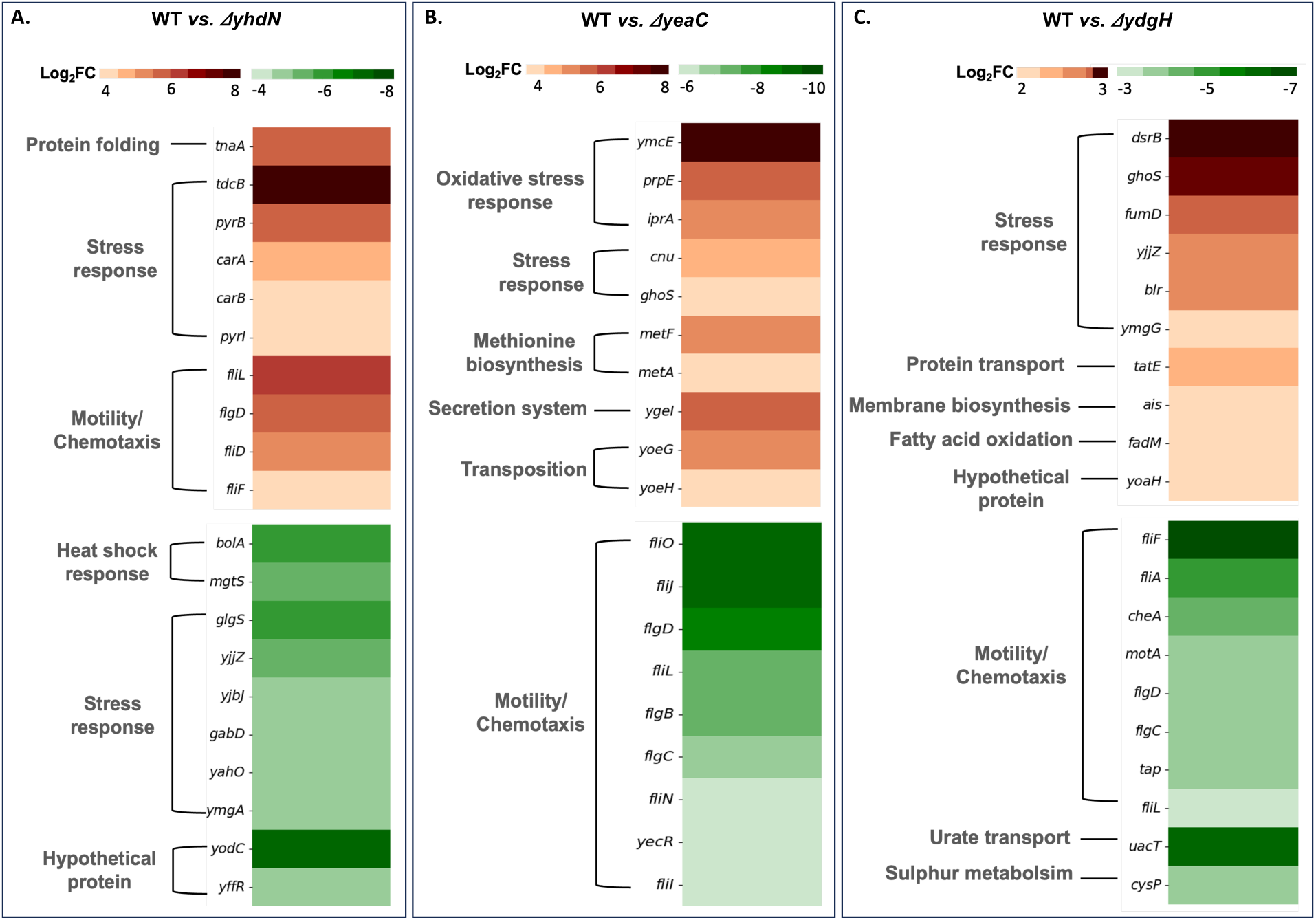
Comparative differential gene expression analysis in A. wild type (WT) *vs. ΔyhdN*, B. WT *vs. ΔyeaC* and C. WT *vs. ΔydgH* strains. Top ten positively and negatively expressed DEGs are shown. Each row represents a gene, and colour intensity represents the Log2 Fold Change (Log2 FC), with red indicating upregulation in WT and green indicating upregulation in the mutant strains. Functional categories were based on information retrieved from Literature and EcoCyc.

In the deletion mutant (**Δ*yhdN*), the top ten negative DEGs included two genes involved in heat shock, six in some type of stress response and interestingly two other were uncharacterized hypothetical proteins. *bolA* (Log_2_ FC=-5.53), a DNA binding transcriptional regulator, plays a role in cellular stress response including heat stress [148] and *mgtS* (Log_2_ FC=-5.01), a small inner membrane protein encoding gene, is known to be expressed upon heat shock [149] (**Figure 4A**). Other upregulated genes in **Δ*yhdN* involved *glgS* (Log_2_ FC=- 5.34), encoding for a surface composition regulator for biofilms and *ymgA* (Log_2_ FC=-4.44) encoding a putative two-component system connector protein, known to play an important role in biofilm formation (EcoCyc). Upregulation of these genes in the deletion mutant suggests a compensatory mechanism to mitigate stress. Additionally, a HP-encoding gene, *yodC* was also upregulated (Log_2_ FC=-6.39) in **Δ*yhdN*. Interestingly, this gene was predicted to play a role in oxidoreduction (**Table1)**. The results of this transcriptomic experiment could further aid in forming a hypothesis of this gene for *in vitro* testing. Another HP-encoding gene *yffR* (Log_2_ FC=-4.85), was predicted to play a role in stress response (this study, Table 1) which aligns with the fact that a majority of stress-response genes were upregulated in the WT, when gene *yhdN* is present.

All observations highlight WT strain’s prioritization of growth and motility, possibly facilitating better environmental sensing and resource acquisition under stress while the absence of *yhdN* gene seems to upregulate other heat shock and stress response genes which reorient metabolic processes towards alternative survival strategies such as biofilm formation. Hence, the protein encoding gene *yhdN* is predicted to be a stress response gene involved in the regulation of heat shock or nutrient limitation response, possibly also playing a broader role in general stress response.

#### WT *vs. ΔyeaC*

In WT strain, the *yeaC* gene exhibited a significant upregulation after the 5h, with a log_2_ FC of 3 relative to the 1h time point, where its expression was undetectable. This log_2_FC corresponds to an 8-fold increase in expression. Hence, the 5h time point was chosen for the analysis. A total of 156 DEGs for WT *vs*. *ΔyeaC* were obtained after filtering (*p_adj_*≤0.1). In WT *vs. yeaC*, 87 genes had a positive Log_2_ FC while 69 had a negative Log_2_ FC.

Analysis of the top ten upregulated DEGs in the WT strain resulted in three oxidative stress response genes, two other stress response genes, two methionine biosynthesis genes and three genes involved in bacterial secretion system and transposition (EcoCyc). The upregulation of the latter three genes in WT was difficult to explain, since there was no conclusive literature available (**Figure 4B**). Significantly upregulated DEGs involved *ymcE* (Log_2_ FC=6.81) which provides tolerance to n-butanol [145], and is also known to provide protection against oxidative stress [150], *prpE* (Log_2_ FC=4.75) which responds to hydrogen peroxide [151] and *iprA* (Log_2_ FC=4.34), which encodes for an inhibitor of hydrogen peroxide [152]. *metF* (Log_2_ FC=4.49), a methionine biosynthesis gene, is also known to provide protection against oxidative stress: Methionine is a precursor in the synthesis of cysteine, which is a component of glutathione—a major intracellular antioxidant. Glutathione is known to neutralize reactive oxygen species [153].

All of the upregulated genes in *ΔyeaC* were involved in motility/chemotaxis-based functions (**Figure 4B**). Significantly upregulated DEGs included genes like *fliF* (Log_2_ FC=-9.86), *fliO* (Log_2_ FC=-9.26) and *fliJ* (Log_2_ FC=-9.26). The upregulation of these genes in *ΔyeaC* signify that the cells prioritize the expression of motility-based genes. They are known to aid the bacterium in adapting to environmental changes by modulating their expression [154]. Moreover, there is often a trade-off between motility and stress resistance. Studies have shown that hypermotile *E. coli* strains, which exhibit increased expression of motility genes, may have reduced expression of stress resistance genes, as was also observed in the downregulated DEGs in the WT. Hence, as opposed to WT, the deletion mutant prioritizes motility and chemotaxis to mitigate oxidative stress.

#### WT *vs. ΔydgH*

The *ydgH* gene demonstrated an upregulation after the 5h, with a log_2_FC of approximately 3.7 as compared to 1 hour, indicating a 13-fold increase in expression. Hence, also in this case the 5h time point was chosen for further analysis. A total of 231 DEGs were obtained after filtering (*p_adj_*≤0.1), resulting in 111 genes with a positive Log_2_ FC and 120 genes with a negative Log_2_ FC.

Top ten positive DEGs (Log_2_ FC≥1.5) involved six stress response genes, one protein transport gene, one membrane biosynthesis gene, one fatty acid oxidation gene and one HP-encoding gene (**Figure 4C**). Significantly upregulated genes were *dsrB* (Log_2_ FC=3.89), encoding for a putative stress response protein [155] and *ghoS,* (Log_2_ FC=3.65), an antitoxin protein which prevents cell death [156]. Gene *fadM* (Log_2_ FC=2.4), encoding a thioesterase, is responsible for breaking down fatty acids into acetyl-CoA units, which can then be utilized in energy production [157]. Another gene, *tatE* (Log_2_ FC=2.69) encoding for a transporter, is involved in transport of folded proteins and maintaining of cellular redox balance [158]. Gene *yoaH,* a HP-encoding gene whose function was previously unclear from **Table 1**, might also play a role in oxidative stress response and should be further investigated. The role of lipopolysaccharide biosynthesis gene *ais* (Log_2_ FC= 2.41) was, however, unclear.

In the deletion mutant (**Δ*ydgH*), the top ten negative DEGs included eight motility/chemotaxis genes along with one sulphur metabolism and one urate transport gene (**Figure 4C**): *fliF* (Log_2_ FC=-6.92) and *fliA* (Log_2_ FC=-5.18) play a crucial role for motility (EcoCyc), *uacT* (Log_2_ FC=- 6.22) is involved in urate transport, which has already been studied to reduce the effects of oxidative stress in *E. coli* [159]. The role of gene *cysP* regarding oxygen stress, which is involved in sulphur metabolism [160] is unclear at the moment and calls for further investigation.

In WT, the upregulation of genes involved in response to stress, i.e. *ghoS* suggest that the cells prioritize conservation of resources by entering a dormant state, i.e. formation of persister cells [156]. Due to the absence of *ΔydgH* the cell seems to attempt to compensate for the loss of oxidoreductive function by utilizing genes related to motility and chemotaxis (**Figure 4C**). This increased expression could be an adaptive response to maintain survival under continuous exposure to oxidative stress. Hence, similar to the previous case (WT *vs. ΔyeaC*), more stress response genes were upregulated in WT while motility/chemotaxis genes were upregulated in *ΔydgH*.

### Conclusion and Outlook

The functional characterization of hypothetical protein (HP)-encoding genes remains a critical challenge, since the rapid accumulation of sequencing data has long surpassed the rate of possible experimental characterization *in vivo* and *in vitro*. While there have been notable advancements in methodologies for protein function prediction in the recent past, several challenges persist, including the heavy reliance on pre-existing data inferred from model organisms. Unfortunately, even in the best studied model organisms a significant amount of protein encoding genes could so far not been annotated.

AI-driven approaches, such as machine learning (ML) algorithms, now offer powerful tools to analyze vast omics datasets. In this study we employed a comprehensive approach, integrating independent component analyses to decipher transcriptional regulatory networks, deep learning for structural homolog prediction, and various other bioinformatic tools to characterize HPs that have no functional information available in public databases. With this methodology putative/probable functions for at least 64% of 95 hypothetical HPs encoding genes from *E. coli* K-12 could be extrapolated, facilitating the subsequent functional validation of these proteins through *in vitro* and *in vivo* experiments in the future. For the remaining 36% HPs, however, more high-quality datasets including metabolomics and proteomics are probably needed, as these genes are often seen to be expressed under very specific environmental conditions from which only one dataset was available. Nevertheless, for most proteins it is possible to derive at least some characteristics from their general sequence properties, whether there is evidence of expression, their possible cellular location, details of homologs, or whether they are predicted or demonstrated to be expressed as part of an operon and/or regulon. Taking these information into account in databases will aid in the improvement of using AI tools for annotation in the future.

In this study we furthermore provided an experimental feedback loop for three HP-encoding genes to verify the *in silico* prediction. Gene *yhdN* ought to play a role in the cells’ responding to heat shock and nutrient limitation, which could be confirmed by comparing the growth curves and transcriptional responses of wild type and *ΔyhdN* deletion mutant strains. For the second candidate *yeaC, in silico* predictions suggested a role in aerobic respiration. Experimental evidence supported this hypothesis by the upregulation of oxidative stress response and methionine biosynthesis genes, hence it is likely involved in scavenging reactive oxygen species. In contrast, the significant upregulation of “Motility”/Chemotaxis” genes in the deletion mutant suggest that during the absence of *yeaC*, bacteria may enhance motility pathways as an adaptive mechanism under oxidative stress. The third candidate *ydgH*, was predicted *in silico* to play a role in oxidoreduction, which could also be confirmed by the *in vivo* experiments showing its role in the protection against oxidative stress. Similar to *yeaC,* in the mutant an upregulation of motility/chemotaxis genes indicate that in the absence of *ydgH*, the cell may compensate for reduced oxidoreductive function. However, it is important to acknowledge that our understanding still remains incomplete since the functional landscape of proteins is vast and complex. Future studies and more omics datasets are needed to further refine and expand upon our findings, exploring the broader implications of these proteins under different environmental conditions.

In conclusion, this study represents a significant step forward in elucidating the functions of previously uncharacterized HPs with no information in existing knowledge databases. Leveraging AI-driven annotations and integrating them with experimental laboratory work will link new functional roles of genes, discover previously unknown cellular and metabolic processes and maybe even biotechnological potentials. In the future this approach might also aid in the isolation of new and so far uncultured microbial species, i.e. microbial dark matter, by harnessing meta-omics datasets and identifying genes critical for growth and adaptation, offering clues to optimize culture conditions.

## Methods

### Identification and enumeration of hypothetical proteins (HPs)

For the comprehensive analysis of *Escherichia coli* K-12 (*E. coli*) substrains MG1655 and BW25113, the complete genome sequences available in the NCBI RefSeq database under the accession number GCF_000005845.2 and GCF_004355105.2 were used, respectively. Ribosomal RNA (rRNA), transfer RNA (tRNA), non-coding RNA (ncRNA) were filtered out. CDS based annotations were used to find hypothetical proteins (HPs). HPs were identified based on keywords like “Putative”, “Putative uncharacterized”, “Uncharacterized protein” and “DUF (domain of unknown function)-domain containing protein”, “UPF” (unknown protein family) as well as proteins having a ‘Protein Y…’ pattern. Further information was extracted from the extensively curated databases EcoCyc version 26.1 [39], RegulonDB version 11.2 [11], EggNOG v6.0 [12] and UniProt release 2023_04 for proteome UP000000625 [1]. In order for a HP to be classified as a protein with no characterization (target group of this study), it had to fulfill all of the following criteria:

1. The **EcoCyc** protein summary showed the statement ‘No information about this’. Additionally, the protein had to be marked as ‘Uncharacterized’. Proteins which had the statement but a ‘Partial’ characterization in EcoCyc were further investigated using UniProt.
2. In **UniProt** the protein lacked any functional information in the ‘Function’ section and was assigned an annotation score of ‘1’, and a ‘protein existence’ score of ‘4’ signifying that the protein, although predicted to be expressed, lacked information based on experimental evidence or orthologs in closely related species. For the “Function” section, a score of ‘1’ is the lowest rating in a five-point-scoring system and for the “protein existing” category a score of ‘4’ out of ‘5’, where ‘5’ indicates a higher level of uncertainty regarding the protein’s expression.
3. In the **RegulonDB** database it had the label “Weak” evidence for an annotation in the evidence section and no additional information in the ‘Note’ subsection.
4. In the **EggNOG** database the protein was either classified under the cluster of orthologous groups (COG) with a functional category ‘S’, where ‘S’ designates proteins belonging to an unknown function category or it lacked an ortholog which was displayed as ‘No orthologs found’.

95 of the 4403 *E. coli* K-12 protein encoding genes fulfilled these criteria. All scripts for the filtering and enumeration of uncharacterized HPs can be found on Figshare (*Notebooks>HP_curation*). In addition, the here identified HP-encoding genes were compared with the ones identified by Ghatak et al. in 2019 [8] to check for their current status.

### Transcriptomic data curation and processing

RNA-sequencing data from NCBI for analyzing the expression patterns of genes was collected for *E. coli* K-12 substrains MG1655 and BW25113 using an adapted workflow for *Bacillus subtilis* [32]. For substrain MG1655 1854 datasets and for substrain BW25113 662 datasets were downloaded and processed. Raw FASTQ files were obtained using sratoolkit 3.0.1 [161], and subsequent read trimming was performed using Trim Galore 0.6.5 [162] with the default settings. Trimmed reads were then subjected to FastQC 0.11.9 [163]. Alignments to the genomes were done using Bowtie 1.2.3 [164]. The read direction was determined using RSeQC 3.0.1 [165] before generating read counts using featureCounts [166]. Finally, all quality control metrics were compiled using MultiQC [167] and the expression compendium was obtained as log-transformed Transcripts per Million (log-TPM) values. Quality metrics were established to filter and ensure the selection of high-quality datasets with the following modifications [168]: Briefly, the datasets were considered to be of high quality if they passed the fastqc parameters ‘per_base_sequence_quality’, ‘per_sequence_quality_scores’, ‘per_base_n_content’, ‘adapter_content’ and had at least 500,000 reads mapped to coding sequences from the reference genome. To preserve biological variability and prevent loss of information, samples with no replicates were also retained in the dataset. Short non-coding transcripts (<100 nucleotides) and extremely low-expression transcripts (Fragments Per Kilobase per Million mapped reads < 10) were removed to reduce noise. 779 datasets for substrain MG1655 and 135 datasets for substrain BW25113 were then used for further analysis. All other information on the *E. coli* K-12 genes including their transcription units, descriptions etc. were sourced from EcoCyc by utilizing the SmartTables feature [39]. Annotations for clusters of orthologous groups (COG) were obtained from EggNOG v6.0 [12] including Gene Ontology [169]. The transcriptional regulatory network (TRN) comprising information on genes and their governing regulators (i.e. transcription factors controlling the expression of genes) were sourced from RegulonDB version 12.0 [11] and EcoCyc [39] as described by Sastry et al. [32] In order to mitigate any batch effects which might be caused due to the usage of diverse datasets, a reference condition (wild type *E. coli* K-12 substrain MG1655 in M9 minimal medium facilitated with glucose and essential micronutrients under non-stressful conditions) was used [30]. This was done by subtracting the mean expression of two replicates of the chosen reference condition from the entire gene expression dataset and calculating the log2-fold-change i.e. relative changes in expression. This normalization method allowed for the comparison of gene expression differences relative to a common baseline as previously demonstrated in Sastry et al. [30]. The normalized data was then used as input for machine learning (*Data processing>Nextflow*).

### Independent Component Analysis of Transcriptomic Data

Independent Component Analysis (ICA) developed by McCon et al. -named OptICA-was applied to prevent both over-decomposition and under-decomposition of transcriptomic datasets [27]. The application of ICA to the normalized *E. coli* K-12 substrain MG1655 RNAseq data resulted in two matrices: the first matrix, denoted as the M matrix, encompassing the robust independent components named iModulons, while the second matrix, termed the A matrix, comprised the corresponding activities of iModulons over different growth conditions. Additionally, a PyModulon package was used in conjunction with Jupyter notebooks to characterize the iModulons [32]. The scripts containing the specific utilization of resources and curation of iModulons can be found on Figshare (*OptICA* and *iModulon curation*). The information from substrain MG1655 was then used to infer the iModulons and their member genes for substrain BW25113, since only for the latter strain single-gene deletion mutants were available for *in vitro* experimentation. The HP-encoding genes, the regulators governing them, and their derived functions can be found in **Table 1**.

Possible functions for the 95 HP-encoding genes were further identified through regulators governing their gene expression, as provided by RegulonDB and EcoCyc and/or by Gene Ontology (GO) terms derived from co-regulating genes using the PANTHER (Protein ANalysis THrough Evolutionary Relationships) tool [33]. Here, member genes of an iModulon (including the gene of interest, i.e. the HP-encoding gene) were used as input to obtain GO terms to see if any were overrepresented. Statistically significant GO categories with the default false Discovery Rate (FDR)-adjusted p-value (<0.05) were manually identified for each HP (*Supplementary Table 1*). These gene-associated functional categories can be found in **Table 1** and **Figure 2**.

### Metadata extraction

Metadata conditions for all datasets were retrieved either manually from the sequence read archive (SRA), *via* a semi-automated approach [167] or by utilizing data from the Gene Expression Omnibus Database using the GEOparse tool [170] to obtain experimental information not available *via* the NCBI’s Entrez Programming Utilities (E-utilities) [42]. As a result, from 779 quality checked datasets for the substrain MG1655, metadata for 602 datasets could be retrieved with confidence and termed as ‘Explicit metadata’. 113 datasets were termed as ‘poor metadata’, meaning that information on one or more experimental conditions (such as media composition, temperature, pH, etc.) was missing or incomplete. Lastly, datasets with ‘unclear metadata’ lacked all information. For the substrain BW25113, 128 datasets were ‘Explicit’ while seven were ‘unclear’ (*Notebooks>metadata_curation*). Additionally, metadata condition for the highest expression of the HP-encoding genes against the reference gene *frr* were extracted from the Tjaden dataset [44] in order to account for the loss of datasets after quality filtering (also see below). The *frr* gene expression is typically consistent across different conditions which was also observed when tested as reference along with other genes like *rpoC*, *rpoD* for the uncharacterized HPs [this study, 171]. Hence, gene *frr* was chosen as the reference gene serving as a control to validate the relative expression levels of other genes. The extensive nature of this dataset ensured a broad representation of gene expression profiles, in addition to the datasets procured for OptICA analysis (*Notebooks> metadata_curation*).

### Co-expression analysis based on public RNA-Seq data

To identify genes with expression patterns similar to those encoding HPs, a Spearman correlation analysis on the Tjaden dataset was performed [172]. This dataset comprises a comprehensive collection of 3.376 *E. coli* strain K-12 RNA-seq datasets, with expression levels quantified in transcripts per million (TPM) [44]. For inferring gene-gene correlations across the transcriptomic datasets for strain *E. coli* K-12, we identified the top ten co-expressed genes to the respective 95 HP-encoding gene and used this information to get an inference on their functions [10] **(Table 1)** (*Notebooks>gene_co-expression*).

### Sub-Cellular Localization

For the purpose of determining the subcellular localizations of the 95 HP-encoding genes, BUSCA (Bologna Unified Subcellular Component Annotator) was employed [173]. This tool integrates various machine learning and bioinformatics tools [174–177] to analyze and recognize patterns in protein sequences that are indicative of specific subcellular localizations. The integrative webserver was used to obtain labels such as ‘Nucleus’, ‘Periplasm/cytoplasm’, ‘Inner/outer membrane’ or ‘extracellular space’. All 95 HP-encoding genes with their predicted sub-cellular localizations can be found in **Table1**.

### Protein-Protein Interactions (PPIs) Analysis

The STRING database (version 12.0) (https://string-db.org) was employed for the identification of PPIs of HP-encoding genes against the proteomic landscape of *E. coli*. The tool incorporates data from diverse origins such as available publications, empirical evidence, predictions derived from co-expression patterns, genomic context analysis, and curated pathway databases. A ‘confidence score’ for each inferred association was derived, quantitatively expressing the robustness of the predicted interaction [178]. In order to keep only the high-scoring associations, a cut-off of 0.7 was utilized [179]. All 95 HP-encoding genes and their interacting partner proteins are listed in **Table 1**.

### Analysis of Gene Context

Shared functional relationships were analyzed using the genomic context of *E. coli* strain K-12 (*Supplementary Figure 1*), particularly the genes located upstream and downstream of the 95 genes of interest. A gene list comprising of gene names was used to create a SmartTable in the EcoCyc database to examine the local genomic neighbourhood and visualize the genes upstream and downstream, including their orientations [36]. (**Table 1)**.

### Detection of Remote Protein Homology

In order to identify distantly homologous sequences, HP sequences were used as input for the HHblits tool (version 3.3.0) available in the MPI Bioinformatics Toolkit using a Hidden Markov Model (HMM) iterative sequence search [180]. The HHblits server (https://toolkit.tuebingen.mpg.de/tools/hhblits) takes a single sequence or a multiple sequence alignment as input and iteratively searches through the selected HMM databases [181,182]. HHblits performs sequence search and remote homology detection with higher sensitivity as compared to gold-standard databases like InterPro [180]. Homologous sequences were then searched against the default UniRef30 database [37]. Sequence based remote homologies identified through the HHblits tool are catalogued in **Table 1**.

### Structure-Based Homology

For identifying structural homologs, we used the AlphaFold Protein Structure Database (AFDB) (https://cluster.foldseek.com/), a specialized computational database that includes structural models for approximately 215 million protein sequences. This platform employs deep learning algorithms to scan and match protein structures, enabling the discovery of structural similarities, even among proteins with low sequence identity [183]. We analyzed clusters to perform comparisons of HPs with all protein structures in the AFDB. UniProt accession IDs were used as inputs to identify the cluster to which the HP belongs. The homology-based function information for each of the 95 HP is listed in **Table 1**. A script to extract the information can be found on Figshare (*Notebooks> HP_info_curation*).

### Bacterial strains and growth conditions

*E. coli* K-12 substrain BW25113 was sourced from Horizon Discovery Ltd., United States. Three single-gene knockout mutants from *E. coli* strain K-12 substrain BW25113 *ΔyhdN, ΔyeaC*, and *ΔydgH* were obtained from the KEIO collection (NBRP, Japan). Glycerol stocks were stored in an ultra-low temperature freezer (Thermo Fisher, United States) at −80°C until further processing. Cells were cultured overnight on agar plates from glycerol stocks and single colonies were picked from the agar plate and then used to Luria Bertani (LB) liquid medium at 37°C [184].

### Media and reagents

*E. coli* strains were cultured in LB broth (10 g/L tryptone, 5 g/L yeast extract, and 10 g/L NaCl; pH 7.0) at 37°C while shaking at 200 rpm [184]. M9 liquid medium for the pre-culture was prepared using 42 mM Na_2_HPO_4_, 22 mM KH_2_PO_4_, 8.5 mM NaCl, 18.7 mM NH_4_Cl, 2 mM MgSO_4_, 0.1 mM CaCl_2_ and 11 mM glucose, supplemented with 0.05% casamino acids to ensure non-limiting conditions. For nitrogen limited M9 medium, a NH_4_Cl concentration of 0.16 mM was used without the addition of casamino acids. For solid medium, 1.5% agar was added. 30% H_2_O_2_ was sterile filtered through a 0.22 µm pore size filter to ensure sterility prior to be used in the oxidative stress experiments. For all experiments with H_2_O_2_ the LB was freshly prepared and kept in the dark until use. All chemicals were sourced from Sigma Chemical Co. (St. Louis, MO, USA).

### Measurement of Bacterial Growth Curves

Bacterial cultures were inoculated at a 1:100 dilution unless stated otherwise. Each experiment included triplicates. All optical density values (OD_600_) were measured *via* a cell density meter (Ultrospec 10, biochrom). OD_600_ was measured every 2h for heat shock in LB and M9 or M9 modified medium and every 1h for oxidative stress experiments. Growth curves were generated for each set of experiments including the wild type and mutant strain in stressed and non-stressed conditions (controls).

#### Heat stress

A single colony (wild type and Δ*yhdN*) was picked from either LB or M9 agar plates and inoculated into 5 mL LB broth for overnight cultivation at 37°C and shaken with 200 rpm. The overnight culture was then used to inoculate fresh LB medium at a 1:100 dilution, which was then cultured for ∼2-3h at 37°C and 200 rpm until an OD_600_ of 0.4 was reached. This was repeated with the M9 liquid medium (preculture), at 37°C to ensure non limiting conditions. For the nitrogen limitation assay, medium was prepared using 0.16mM NH_4_Cl. To study the effects of heat shock on bacterial growth, the culture was grown to mid-log phase (OD_600_ 0.4), diluted to OD_600_ 0.04 and then heat shocked for 7 minutes in a shaker-incubator at 50°C. After treatment, the bacterial suspension was incubated for 12 h at 37°C. Samples were measured every 2h for 12h.

#### Oxidative stress

For studying the effects of oxidative stress, a single colony (wild type, *ΔyeaC* and *ΔydgH*) was picked from the LB agar plate and inoculated into 5 mL LB for overnight cultivation at 37°C and shaken with 200 rpm. The overnight culture was then used to inoculate fresh LB (dilution of 1:100) and cultured at 37°C and 200 rpm until an OD_600_ of 0.2 was reached. An OD of 0.2 was used since at this stage the cells were actively dividing, making them susceptible to the effects of oxidative stress, allowing for better characterization and analysis of the stress response[137]. H_2_O_2_ was added at a final concentration of 2.5 mM. After treatment, the bacterial suspension was incubated for 5h at 37°C and the OD_600_ was measured hourly.

### RNA Sequencing and data processing

2 ml of cell suspension from the heat shock samples was collected after 2h and 6h of growth. For the oxidative stress experiments, samples were collected after 1h and 5h of growth. The medium was discarded, cell pellets were washed with Phosphate Buffer Saline and mixed with DNA/RNA shield™ (Zymo Research Corp., Freiburg, Germany). All samples were processed according to the manufacturer’s instructions. Samples were preserved at -20°C until RNA extraction. RNA was extracted using the Direct-zol RNA Miniprep Plus Kit (Zymo Research Corp., Freiburg, Germany) according to the manufacturer’s instructions. Total RNA was eluted from columns with 100 μL nuclease-free water and quantified using the Qubit RNA Broad Range Assay Kit (Thermo Fisher Scientific Inc., Waltham, MA, USA). The quality was assessed by running an RNA-nano chip on an Agilent bioanalyzer (Agilent Technologies, Waldbronn, Germany). Samples with an RNA integrity score ≥8 were used for further processing [185]. The rRNA was removed from total RNA preparations using the NEBNext rRNA Depletion (Bacteria) kit (New England Biolabs, Frankfurt, Germany). Paired-end library preparation was done using the NEBNext Ultra II Directional RNA Library Prep Kit for Illumina (New England Biolabs, Frankfurt, Germany) following the manufacturer’s instructions with an average insert size of 200 bp. Libraries were run on a NextSeq 550™ (Illumina, San Diego, CA, United States).

The RNAflow pipeline [186] was implemented using a conda environment. Sequence reads were checked for quality using FastQC v0.11.9 [187]. Raw reads were processed using fastp v0.20.0 [188], which employs a sliding window approach for quality trimming and adapter clipping, with the settings: a -5 to -3 cut for front and tail, a default window of 4 and mean quality threshold of 20. Trimmed reads shorter than 15 bps were discarded. The remaining reads were mapped onto the *E. coli* K-12 reference genome (NZ_CP009273.1) using HISAT v2.1.0 [189]. The generated output was converted into a BAM format using samtools-1.20 [190], followed by the processes of sorting and indexing. A gene transfer format (GTF) annotation file for *E. coli* substrain BW25113 was used to filter for gene and pseudogene features. FeatureCounts v2.0.1 [191] was used to quantify the mapped reads based on the gene annotations. Processed datasets were summarized in a MultiQC v1.9 [167] generated file. were normalized by the DESeq2 module (v1.28.0) in R (version 4.3.2) using a variance stabilizing transformation [192].

### Comparative gene expression analysis of wild type and isogenic mutant strains

Sequencing reads were obtained from the transcriptomic datasets of wild-type (WT), *ΔyhdN*, *ΔyeaC,* and *ΔydgH* mutants, respectively. Non-stressed control and heat shocked samples from WT and *ΔyhdN* at the 6h timepoint and non-stressed control and oxidative stress samples from WT, *ΔyeaC,* and *ΔydgH* at the 5h timepoint were used to identify up- or downregulated genes. DESeq2 [192] was utilized to examine expression differences of genes in the WT and mutant strains under both control and stress conditions. To focus on the specific effect of stress on gene expression rather than the difference of WT *vs.* deletion mutants, read counts under control conditions were subtracted from the read counts under stress condition [193]. Genes with an adjusted P-value (Padj) of 0.1 or less were classified as differentially expressed to avoid false positives. A cut-off log_2_ fold change (log_2_FC) of 1.5 was chosen for further analysis [194]. A log_2_FC of 1.5 means that a gene was 2.83 times more expressed in one strain *vs.* the other, a log_2_FC of 2 means that a gene was 4 times more expressed and so on. Significant positive log_2_FC indicated genes that were upregulated in WT relative to the mutant, while negative values showed an upregulation in the mutant *vs*. WT. To determine the functional categories of these genes, the EcoCyc database was used [36]. Information of the differentially expressed genes can be found on Figshare (*DEGs>Table1*).

## Data availability

All code and scripts used in this study are available on Figshare. These resources can be accessed at https://figshare.com/s/0ede175c510cf201e7c2.

Raw sequencing data that supports the findings of this study have been deposited in the Sequence Read Archive with the BioProject ID: PRJNA1209969.

## Competing interests

The authors declare that they have no conflict of interest.

## Funding

This work was supported by the Karlsruhe Institute of Technology and the Helmholtz Society [POF4; 5207.0004.0012].

## Authors’ contributions

S.C, H.A. and Z.A did the data analyses, S.C. did the *in vitro* experiments, S.C and A.K.K. wrote the manuscript, A.K.K. provided the funding.

## Acknowledgements

The authors acknowledge the support by the state of Baden-Württemberg through bwHPC. We thank Dr. Gunnar Sturm for technical support and insightful discussions and David Thiele for library preparation assistance.

## Notes

https://figshare.com/s/0ede175c510cf201e7c2

### Competing Interest Statement

The authors have declared no competing interest.

### Summary of Updates

This version of the manuscript has been revised to update the title and the abstract. The scientific content of the manuscript remains unchanged. No changes were made to the author list, figures, or supplementary materials.

## References

1. Consortium TU. UniProt: the Universal Protein Knowledgebase in 2023. Nucleic Acids Res. 2023;51:D523–31; doi:10.1093/nar/gkac1052

2. Paysan-Lafosse T, Blum M, Chuguransky S, Grego T, Pinto BL, Salazar GA, et al. InterPro in 2022. Nucleic Acids Res. 2023;51:D418–27; doi:10.1093/nar/gkac993

3. Mistry J, Chuguransky S, Williams L, Qureshi M, Salazar GA, Sonnhammer ELL, et al. Pfam: The protein families database in 2021. Nucleic Acids Res. 2021;49:D412–9; doi:10.1093/nar/gkaa913

4. Sigrist CJA, de Castro E, Cerutti L, Cuche BA, Hulo N, Bridge A, et al. New and continuing developments at PROSITE. Nucleic Acids Res. 2013;41:D344–7; doi:10.1093/nar/gks1067

5. Letunic I, Khedkar S, Bork P. SMART: recent updates, new developments and status in 2020. Nucleic Acids Res. 2021;49:D458–60; doi:10.1093/nar/gkaa937

6. Lu S, Wang J, Chitsaz F, Derbyshire MK, Geer RC, Gonzales NR, et al. CDD/SPARCLE: the conserved domain database in 2020. Nucleic Acids Res. 2020;48:D265–8; doi:10.1093/nar/gkz991

7. Durairaj J, Waterhouse AM, Mets T, Brodiazhenko T, Abdullah M, Studer G, et al. Uncovering new families and folds in the natural protein universe. Nature. 2023;622:646–53; doi:10.1038/s41586-023-06622-3

8. Ghatak S, King ZA, Sastry A, Palsson BO. The y-ome defines the 35% of *Escherichia coli* genes that lack experimental evidence of function. Nucleic Acids Res. 2019;47:2446–54; doi:10.1093/nar/gkz030

9. Ardern Z, Chakraborty S, Lenk F, Kaster A-K. Elucidating the functional roles of prokaryotic proteins using big data and artificial intelligence. FEMS Microbiol Rev. 2023;47; doi:10.1093/femsre/fuad003

10. Keseler IM, Gama-Castro S, Mackie A, Billington R, Bonavides-Martínez C, Caspi R, et al. The EcoCyc Database in 2021. Front Microbiol. 2021;12; doi:10.3389/fmicb.2021.711077

11. Salgado H, Gama-Castro S, Lara P, Mejia-Almonte C, Alarcón-Carranza G, López-Almazo AG, et al. RegulonDB v12.0: a comprehensive resource of transcriptional regulation in *E. coli* K-12. Nucleic Acids Res. 2024;52:D255–64; doi:10.1093/nar/gkad1072

12. Hernández-Plaza A, Szklarczyk D, Botas J, Cantalapiedra CP, Giner-Lamia J, Mende DR, et al. eggNOG 6.0: enabling comparative genomics across 12 535 organisms. Nucleic Acids Res. 2023;51:D389–94; doi:10.1093/nar/gkac1022

13. Mills CL, Beuning PJ, Ondrechen MJ. Biochemical functional predictions for protein structures of unknown or uncertain function. Comput Struct Biotechnol J. 2015;13:182–91; doi:10.1016/j.csbj.2015.02.003

14. Zhang C, Freddolino PL, Zhang Y. COFACTOR: improved protein function prediction by combining structure, sequence and protein–protein interaction information. Nucleic Acids Res. 2017;45:W291–9; doi:10.1093/nar/gkx366

15. Makrodimitris S, van Ham RCHJ, Reinders MJT. Improving protein function prediction using protein sequence and GO-term similarities. Bioinformatics. 2019;35:1116–24. Available from: doi:10.1093/bioinformatics/bty751

16. Saha S, Chatterjee P, Basu S, Nasipuri M, Plewczynski D. FunPred 3.0: improved protein function prediction using protein interaction network. PeerJ. 2019;7. doi: 10.7717/peerj.6830

17. Varadi M, Tsenkov M, Velankar S. Challenges in bridging the gap between protein structure prediction and functional interpretation. Proteins: Structure, Function, and Bioinformatics. 2023; doi:10.1002/prot.26614

18. Vincent AT. Bacterial hypothetical proteins may be of functional interest. Frontiers in Bacteriology. 2024;3; doi:10.3389/fbrio.2024.1334712

19. Jeffery CJ. Current successes and remaining challenges in protein function prediction. Frontiers in Bioinformatics. 2023;3; doi:10.3389/fbinf.2023.1222182

20. Pavlopoulos GA, Baltoumas FA, Liu S, Selvitopi O, Camargo AP, Nayfach S, et al. Unraveling the functional dark matter through global metagenomics. Nature. 2023;622:594– 602; doi:10.1038/s41586-023-06583-7

21. Escudeiro P, Henry CS, Dias RPM. Functional characterization of prokaryotic dark matter: the road so far and what lies ahead. Curr Res Microb Sci. 2022;3:100159; doi: 10.1016/j.crmicr.2022.100159

22. da Costa WLO, Araújo CL de A, Dias LM, Pereira LC de S, Alves JTC, Araújo FA, et al. Functional annotation of hypothetical proteins from the *Exiguobacterium antarcticum* strain B7 reveals proteins involved in adaptation to extreme environments, including high arsenic resistance. PLoS One. 2018;13:e0198965; doi:10.1371/journal.pone.0198965

23. Grünberger F, Knüppel R, Jüttner M, Fenk M, Borst A, Reichelt R, et al. Exploring prokaryotic transcription, operon structures, rRNA maturation and modifications using Nanopore-based native RNA sequencing. bioRxiv. 2020;2019.12.18.880849; doi: 10.1101/2019.12.18.880849

24. Dall’Alba G, Casa PL, Abreu FP de, Notari DL, de Avila e Silva S. A Survey of Biological Data in a Big Data Perspective. Big Data. 2022;10:279–97; doi:10.1089/big.2020.0383

25. Chen J, Gu Z, Lai L, Pei J. In silico protein function prediction: the rise of machine learning-based approaches. 2023;3:487–510; doi:10.1515/mr-2023-0038

26. Alteri CJ, Lindner JR, Reiss DJ, Smith SN, Mobley HLT. The broadly conserved regulator PhoP links pathogen virulence and membrane potential in *Escherichia coli*. Mol Microbiol. 2011;82:145–63; doi:10.1111/j.1365-2958.2011.07804.x

27. Mcconn JL, Lamoureux CR, Poudel S, Palsson BO, Sastry A V. Optimal dimensionality selection for independent component analysis of transcriptomic data; doi:10.1186/s12859-021-04497-7

28. Yu J, Li M, Wang J, Hamushan M, Jiang F, Wang B, et al. Identification of *Staphylococcus aureus* virulence-modulating RNA from transcriptomics data with machine learning. Virulence. 2023;14:2228657; doi:10.1080/21505594.2023.2228657

29. Comon P. Independent component analysis, A new concept? Signal Processing. 1994;36:287–314; doi:10.1016/0165-1684(94)90029-9

30. Sastry A V, Gao Y, Szubin R, Hefner Y, Xu S, Kim D, et al. The *Escherichia coli* transcriptome mostly consists of independently regulated modules. Nat Commun. 2019;10:5536; doi:10.1038/s41467-019-13483-w

31. Sastry A V, Poudel S, Rychel K, Yoo R, Lamoureux CR, Chauhan S, et al. Title Mining all publicly available expression data to compute dynamic microbial transcriptional regulatory networks Authors; doi:10.1101/2021.07.01.450581

32. Rychel K, Sastry A V., Palsson BO. Machine learning uncovers independently regulated modules in the *Bacillus subtilis* transcriptome. Nat Commun. 2020;11. doi: 10.1038/s41467-020-20153-9

33. Thomas PD, Ebert D, Muruganujan A, Mushayahama T, Albou L-P, Mi H. PANTHER: Making genome-scale phylogenetics accessible to all. Protein Science. 2022;31:8–22; doi:10.1002/pro.4218

34. Savojardo C, Martelli PL, Fariselli P, Profiti G, Casadio R. BUSCA: an integrative web server to predict subcellular localization of proteins. Nucleic Acids Res. 2018;46:W459–66; doi:10.1093/nar/gky320

35. Szklarczyk D, Kirsch R, Koutrouli M, Nastou K, Mehryary F, Hachilif R, et al. The STRING database in 2023: protein–protein association networks and functional enrichment analyses for any sequenced genome of interest. Nucleic Acids Res. 2023;51:D638–46; doi: 10.1093/nar/gkac1000

36. D KP, Suzanne P, Ron C, Anamika K, Markus K, E MP, et al. The EcoCyc Database (2023). EcoSal Plus. 2023;11:eesp-0002-2023; doi:10.1128/ecosalplus.esp-0002-2023

37. Gabler F, Nam S-Z, Till S, Mirdita M, Steinegger M, Söding J, et al. Protein Sequence Analysis Using the MPI Bioinformatics Toolkit. Curr Protoc Bioinformatics. 2020;72:e108; doi:10.1002/cpbi.108

38. Barrio-Hernandez I, Yeo J, Jänes J, Mirdita M, Gilchrist CLM, Wein T, et al. Clustering predicted structures at the scale of the known protein universe. Nature. 2023;622:637–45; doi:10.1038/s41586-023-06510-w

39. Karp PD, Billington R, Caspi R, Fulcher CA, Latendresse M, Kothari A, et al. The BioCyc collection of microbial genomes and metabolic pathways. Brief Bioinform. 2019;20:1085–93; doi:10.1093/bib/bbx085

40. Choudhary S. pysradb: A Python package to query next-generation sequencing metadata and data from NCBI Sequence Read Archive. F1000Res. 2019;8:532; doi: 10.12688/f1000research.18676.1

41. Gumienny R. GEOparse. Rev.0a257463. 2015. Available from: https://geoparse.readthedocs.io/en/latest/

42. Liu M, Durfee T, Cabrera JE, Zhao K, Jin DJ, Blattner FR. Global transcriptional programs reveal a carbon source foraging strategy by *Escherichia coli*. J Biol Chem. 2005;280:15921– 7; doi: 10.1074/jbc.m414050200

43. Tjaden B. *Escherichia coli* transcriptome assembly from a compendium of RNA-seq data sets. RNA Biol. 2023;20:77–84; doi:10.1080/15476286.2023.2189331

44. Rawangkan A, Yosboonruang A, Kiddee A, Siriphap A, Pook-In G, Praphasawat R, et al. Restoring Ampicillin Sensitivity in Multidrug-Resistant *Escherichia coli* Following Treatment in Combination with Coffee Pulp Extracts. J Microbiol Biotechnol. 2023;33:1–10; doi:10.4014/jmb.2304.04051

45. Manat G, El Ghachi M, Auger R, Baouche K, Olatunji S, Kerff F, et al. Membrane Topology and Biochemical Characterization of the *Escherichia coli* BacA Undecaprenyl-Pyrophosphate Phosphatase. PLoS One. 2015;10:e0142870; doi:10.1371/journal.pone.0142870

46. Hajra D, Nair AV, Chakravortty D. An elegant nano-injection machinery for sabotaging the host: Role of Type III secretion system in virulence of different human and animal pathogenic bacteria. Phys Life Rev. 2021;38:25–54; doi:10.1016/j.plrev.2021.05.007

47. Prakash A, Dutta D. Bicyclomycin generates ROS and blocks cell division in *Escherichia coli*. PLoS One. 2024;19:e0293858; doi:10.1371/journal.pone.0293858

48. Ogawa A, Kojima F, Miyake Y, Yoshimura M, Ishijima N, Iyoda S, et al. Regulation of constant cell elongation and Sfm pili synthesis in *Escherichia coli* via two active forms of FimZ orphan response regulator. Genes to Cells. 2022;27:657–74; doi:10.1111/gtc.12982

49. Lamark T, Røkenes TP, McDougall J, Strøm AR. The complex bet promoters of *Escherichia coli*: regulation by oxygen (ArcA), choline (BetI), and osmotic stress. J Bacteriol. 1996;178(6):1655–62; doi:10.1128/jb.178.6.1655-1662.1996.

50. Kambampati R, Lauhon CT. Evidence for the Transfer of Sulfane Sulfur from IscS to ThiI during the in Vitro Biosynthesis of 4-Thiouridine in *Escherichia coli* tRNA*. Journal of Biological Chemistry. 2000;275:10727–30; doi:10.1074/jbc.275.15.10727

51. Laura F-G, Xinyu G, Joy K, Sooyeon S, E BM, Rodolfo G-C, et al. Single-cell analysis reveals that cryptic prophage protease LfgB protects *Escherichia coli* during oxidative stress by cleaving antitoxin MqsA. Microbiol Spectr. 2024;12:e03471–23; doi:10.1128/spectrum.03471-23

52. Zorraquino-Salvo V, Quinones-Soto S, Kim M, Rai N, Tagkopoulos I. Deciphering the genetic and transcriptional basis of cross-stress responses in *Escherichia coli* under complex evolutionary scenarios. bioRxiv. 2014;010595; doi:

53. May T, Okabe S. Enterobactin is required for biofilm development in reduced-genome *Escherichia coli*. Environ Microbiol. 2011;13:3149–62; doi:10.1111/j.1462-2920.2011.02607.x

54. Saier Jr MH, Reddy VS, Tamang DG, Västermark Å. The Transporter Classification Database. Nucleic Acids Res. 2014;42:D251–8; doi:10.1093/nar/gkt1097

55. Muto A, Goto S, Kurita D, Ushida C, Soma A, Himeno H. A leaderless mRNA including tRNA-like sequence encodes a small peptide that regulates the expression of GcvB small RNA in *Escherichia coli*. The Journal of Biochemistry. 2022;171:459–65; doi:10.1093/jb/mvac007

56. Njenga R, Boele J, Öztürk Y, Koch H-G. Coping with stress: How bacteria fine-tune protein synthesis and protein transport. Journal of Biological Chemistry. 2023;299:105163; doi: 10.1016/j.jbc.2023.105163

57. Rodríguez-Ochoa JI, Fragoso-Jiménez JC, Flores N, et al. Global transcriptomic response of *Escherichia coli* to p-coumaric acid. Microb Cell Fact. 2022;21:148; doi:10.1186/s12934-022-01874-6.

58. Bak G, Lee J, Suk S, Kim D, Lee J, Kim K, et al. Identification of novel sRNAs involved in biofilm formation, motility, and fimbriae formation in *Escherichia coli*. Sci Rep. 2015;5:15287; doi:10.1038/srep15287

59. Duarte-Velázquez I, Mora J, Ramírez-Prado J, Aguillón-Bárcenas A, Tornero-Gutiérrez F, Cordero-Loreto E, et al. *Escherichia coli* transcription factors of unknown function: sequence features and possible evolutionary relationships. PeerJ. 2022;10:e13772; doi:10.7717/peerj.13772

60. Saier Jr MH, Reddy VS, Tsu B V, Ahmed MS, Li C, Moreno-Hagelsieb G. The Transporter Classification Database (TCDB): recent advances. Nucleic Acids Res. 2016;44:D372–9; doi:10.1093/nar/gkv1103

61. Su C-C, Rutherford DJ, Yu EW. Characterization of the multidrug efflux regulator AcrR from *Escherichia coli*. Biochem Biophys Res Commun. 2007;361:85–90; doi:10.1016/j.bbrc.2007.06.175

62. Börner J, Friedrich T, Bartkuhn M, Klug G. Ribonuclease E strongly impacts bacterial adaptation to different growth conditions. RNA Biol. 2023;20:120–35; doi:10.1080/15476286.2023.2195733

63. A JN, Melissa F, E FS. Disruption of trehalose periplasmic recycling dysregulates cAMP-CRP signaling in *Escherichia coli* during stationary phase. J Bacteriol. 2023;205:e00292–23; doi:10.1128/jb.00292-23

64. Wang M, Fang K, Hong SMC, Kim I, Jang I-S, Hong SH. Medium chain unsaturated fatty acid ethyl esters inhibit persister formation of *Escherichia coli* via antitoxin HipB. Appl Microbiol Biotechnol. 2018;102:8511–24; doi:10.1007/s00253-018-9271-3

65. Muenter MM, Aiken A, Akanji JO, Baig S, Bellou S, Carlson A, et al. The response of *Escherichia coli* to the alkylating agents chloroacetaldehyde and styrene oxide. Mutation Research/Genetic Toxicology and Environmental Mutagenesis. 2019;840:1–10; doi:10.1016/j.mrgentox.2019.02.001

66. Hideji Y, Yasushi M, Shou F, Akiko S, Masami U, Akira W. YqjD Is an Inner Membrane Protein Associated with Stationary-Phase Ribosomes in *Escherichia coli*. J Bacteriol. 2012;194:4178–83; doi:10.1128/jb.00396-12

67. Hemm MR, Paul BJ, Schneider TD, Storz G, Rudd KE. Small membrane proteins found by comparative genomics and ribosome binding site models. Mol Microbiol. 2008;70:1487– 501; doi:10.1111/j.1365-2958.2008.06495.x

68. Skjold AC, Ezekiel DH. Analysis of lambda insertions in the fucose utilization region of *Escherichia coli* K-12: use of lambda *fuc* and lambda *argA* transducing bacteriophages to partially order the fucose utilization genes. J Bacteriol. 1982;152:120–5<; doi:10.1016/j.mrgentox.2019.02.001>

69. Manuel C, Teresa ZM, Blas B, Gonzalo D-R, F JJ, Andrés VJ, et al. Anaerobic Catabolism of Aromatic Compounds: a Genetic and Genomic View. Microbiology and Molecular Biology Reviews. 2009;73:71–133; 10.1128/mmbr.00021-08

70. Arnim W, A KS, Kirsten J. Time-Dependent Proteome Alterations under Osmotic Stress during Aerobic and Anaerobic Growth in *Escherichia coli*. J Bacteriol. 2006;188:7165–75; doi:10.1128/jb.00508-06

71. Teufel R, Mascaraque V, Ismail W, Voss M, Perera J, Eisenreich W, et al. Bacterial phenylalanine and phenylacetate catabolic pathway revealed. Proc Natl Acad Sci U S A. 2010;107(32):14390–5; doi: 10.1073/pnas.1005399107.

72. Tenorio E, Saeki T, Fujita K, Kitakawa M, Baba T, Mori H, et al. Systematic characterization of *Escherichia coli* genes/ORFs affecting biofilm formation. FEMS Microbiol Lett. 2003;225(1):107–14; doi:10.1016/S0378-1097(03)00507-X.

73. Rodríguez-Montelongo L, Volentini SI, Farías RN, Massa EM, Rapisarda VA. The Cu(II)-reductase NADH dehydrogenase-2 of *Escherichia coli* improves the bacterial growth in extreme copper concentrations and increases the resistance to the damage caused by copper and hydroperoxide. Arch Biochem Biophys. 2006;451:1–7; doi:10.1016/j.abb.2006.04.019

74. Harwani D. Regulation of Gene Expression: Cryptic β-glucoside (bgl) Operon of *Escherichia coli* as a Paradigm. Brazilian Journal of Microbiology. 2014; doi:10.1590/S1517-83822014000400003

75. Anne P, K BI, Aurora L-N, F GJ, Valérie de C-L, D HA. FolX and FolM Are Essential for Tetrahydromonapterin Synthesis in *Escherichia coli* and Pseudomonas aeruginosa. J Bacteriol. 2010;192:475–82; doi:10.1128/jb.01198-09

76. Kotaka Y, Hashimoto M, Lee K, Kato J. Mutations identified in engineered *Escherichia coli* with a reduced genome. Front Microbiol. 2023;14; doi:10.3389/fmicb.2023.1189877

77. Doukyu N, Taguchi K. Involvement of catalase and superoxide dismutase in hydrophobic organic solvent tolerance of *Escherichia coli*. AMB Express. 2021;11:97; doi:10.1186/s13568-021-01258-w

78. Pinske C. The Ferredoxin-Like Proteins HydN and YsaA Enhance Redox Dye-Linked Activity of the Formate Dehydrogenase H Component of the Formate Hydrogenlyase Complex. Front Microbiol. 2018;9; doi:10.3389/fmicb.2018.01238

79. Panta PR, Doerrler WT. A link between pH homeostasis and colistin resistance in bacteria. Sci Rep. 2021;11:13230; doi:10.1038/s41598-021-92718-7

80. Vasilyev N, Liu MMJ, Epshtein V, Shamovsky I, Nudler E. General Transcription Factor from *E. coli* with a Distinct Mechanism of Action. bioRxiv. 2023;2023.09.17.558134; doi: 10.1101/2023.09.17.558134

81. Nasreen M, Purushothaman Nair R, Mcewan A, Kappler U. The Peptide Methionine Sulfoxide Reductase (MsrAB) of Haemophilus influenzae Repairs Oxidatively Damaged Outer Membrane and Periplasmic Proteins Involved in Nutrient Acquisition and Virulence. Antioxidants. 2022;11:1557; doi:10.3390/antiox11081557

82. Yuan J, Jin F, Glatter T, Sourjik V. Osmosensing by the bacterial PhoQ/PhoP two-component system. Proceedings of the National Academy of Sciences. 2017;114:E10792–8; doi:10.1073/pnas.1717272114

83. Steinsiek S, Stagge S, Bettenbrock K. Analysis of *Escherichia coli* mutants with a linear respiratory chain. PLoS One. 2014;9(1):e87307; doi:10.1371/journal.pone.0087307.

84. Akinori K, Hiroyuki T, Ryutaro U. Molecular Characterization of the PhoP-PhoQ Two-Component System in *Escherichia coli* K-12: Identification of Extracellular Mg2+-Responsive Promoters. J Bacteriol. 1999;181:5516–20; doi:10.1128/jb.181.17.5516-5520.1999

85. Podlesek Z, Žgur Bertok D. The DNA Damage Inducible SOS Response Is a Key Player in the Generation of Bacterial Persister Cells and Population Wide Tolerance. Front Microbiol.2020;11; doi:10.3389/fmicb.2020.01785

86. Bitoun JP, Wu G, Ding H. *Escherichia coli* FtnA acts as an iron buffer for re-assembly of iron–sulfur clusters in response to hydrogen peroxide stress. BioMetals. 2008;21:693–703; doi:10.1007/s10534-008-9154-7

87. C BD, J TR. Small Multidrug Resistance Protein EmrE Reduces Host pH and Osmotic Tolerance to Metabolic Quaternary Cation Osmoprotectants. J Bacteriol. 2012;194:5941–8; doi:10.1128/jb.00666-12

88. M FE, E EK, N GL, R GE, M MJ, C MA, et al. YfdW and YfdU Are Required for Oxalate-Induced Acid Tolerance in *Escherichia coli* K-12. J Bacteriol. 2013;195:1446–55; doi:10.1128/jb.01936-12

89. Ralph ET, Guest JR, Green J. Altering the anaerobic transcription factor FNR confers a hemolytic phenotype on *Escherichia coli* K12. Proceedings of the National Academy of Sciences. 1998;95:10449–52; doi:10.1073/pnas.95.18.10449

90. Wang X, Kim Y, Ma Q, Hong SH, Pokusaeva K, Sturino JM, et al. Cryptic prophages help bacteria cope with adverse environments. Nat Commun. 2010;1:147; doi:10.1038/ncomms1146

91. Barabote RD, Rendulic S, Schuster SC, Saier MH. Comprehensive analysis of transport proteins encoded within the genome of *Bdellovibrio bacteriovorus*. Genomics. 2007;90:424–46; doi:10.1016/j.ygeno.2007.06.002

92. Wang J, Ma W, Wang X. Insights into the structure of *Escherichia coli* outer membrane as the target for engineering microbial cell factories. Microb Cell Fact. 2021;20:73; doi:10.1186/s12934-021-01565-8

93. Gurmu D, Lu J, Johnson KA, Nordlund P, Holmgren A, Erlandsen H. The crystal structure of the protein YhaK from *Escherichia coli* reveals a new subclass of redox sensitive enterobacterial bicupins. Proteins: Structure, Function, and Bioinformatics. 2009;74:18–31; doi:10.1002/prot.22128

94. Outten CE, Outten FW, O’Halloran T V. DNA Distortion Mechanism for Transcriptional Activation by ZntR, a Zn(II)-responsive MerR Homologue in *Escherichia coli*. Journal of Biological Chemistry. 1999;274:37517–24; doi:10.1074/jbc.274.53.37517

95. Akbar S, Bhakta S, Sengupta J. Structural insights into the interplay of protein biogenesis factors with the 70S ribosome. Structure. 2021;29:755–767.e4; doi:10.1016/j.str.2021.03.005

96. K VDT, J TL, A CK, L SP, Sima SF. Characterization of the *Escherichia coli* AaeAB Efflux Pump: a Metabolic Relief Valve? J Bacteriol. 2004;186:7196–204; doi:10.1128/jb.186.21.7196-7204.2004

97. Bak G, Lee J, Suk S, Kim D, Young Lee J, Kim K, et al. Identification of novel sRNAs involved in biofilm formation, motility and fimbriae formation in *Escherichia coli*. Sci Rep. 2015;5:15287; doi:10.1038/srep15287

98. P DM, Chi-Fang W, Liang W, J VJ, E BW. DNA Microarray-Based Identification of Genes Controlled by Autoinducer 2-Stimulated Quorum Sensing in *Escherichia coli*. J Bacteriol. 2001;183:5239–47; doi:10.1128/jb.183.18.5239-5247.2001

99. Filenko N, Spiro S, Browning DF, Squire D, Overton TW, Cole J, Constantinidou C. The NsrR regulon of *Escherichia coli* K-12 includes genes encoding the hybrid cluster protein and the periplasmic, respiratory nitrite reductase. J Bacteriol. 2007;189:1203–9; doi:10.1128/jb.00080-07.

100. Khil PP, Camerini-Otero RD. Over 1000 genes are involved in the DNA damage response of *Escherichia coli*. Mol Microbiol. 2002;44:89–105; doi:10.1046/j.1365-2958.2002.02878.x

101. Erol I, Jeong K-C, Baumler DJ, Vykhodets B, Ho Choi S, Kaspar CW. H-NS controls metabolism and stress tolerance in *Escherichia coli* O157:H7 that influence mouse passage. BMC Microbiol. 2006;6:72; doi:10.1186/1471-2180-6-72

102. Zheng L. Catalytic mechanism and function evolvement studies of phosphatases within the haloacid dehalogenase superfamily (HADSF) [dissertation]. Albuquerque (NM): University of New Mexico; 2013. Available from: https://digitalrepository.unm.edu/chem_etds/31.

103. Born TL, Blanchard JS. Structure/function studies on enzymes in the diaminopimelate pathway of bacterial cell wall biosynthesis. Curr Opin Chem Biol. 1999;3:607–13; doi:10.1016/S1367-5931(99)00016-2

104. Hor L, Dobson RCJ, Downton MT, Wagner J, Hutton CA, Perugini MA. Dimerization of Bacterial Diaminopimelate Epimerase Is Essential for Catalysis. J Biol Chem. 2013;288:9238–48

105. Iyer LM, Leipe DD, Koonin E V, Aravind L. Evolutionary history and higher order classification of AAA+ ATPases. J Struct Biol. 2004;146:11–31; doi:

106. Luttringer F, Mulliez E, Dublet B, Lemaire D, Fontecave M. The Zn center of the anaerobic ribonucleotide reductase from *E. coli*. JBIC Journal of Biological Inorganic Chemistry. 2009;14:923–33; doi:10.1007/s00775-009-0505-9

107. Panta PR, Doerrler WT. A link between pH homeostasis and colistin resistance in bacteria. Sci Rep. 2021;11:13230; doi:10.1038/s41598-021-92718-7

108. Dapa T, Fleurier S, Bredeche M-F, Matic I. The SOS and RpoS Regulons Contribute to Bacterial Cell Robustness to Genotoxic Stress by Synergistically Regulating DNA Polymerase Pol II. Genetics. 2017;206:1349–60; doi:10.1534/genetics.116.199471

109. Passariello C, Forleo C, Micheli V, Schippa S, Leone R, Mangani S, et al. Biochemical characterization of the class B acid phosphatase (AphA) of *Escherichia coli* MG1655. Biochimica et Biophysica Acta (BBA) - Proteins and Proteomics. 2006;1764:13–9; doi:10.1016/j.bbapap.2005.08.028

110. Chen SH, Byrne RT, Wood EA, Cox MM. *Escherichia coli rad* gene: a novel function involved in radiation resistance and double-strand break repair. Mol Microbiol. 2015;95:754– 68; doi:10.1111/mmi.12885

111. Chuang SE, Blattner FR. Characterization of twenty-six new heat shock genes of *Escherichia coli*. J Bacteriol. 1993;175:5242–52; doi:10.1128/jb.175.16.5242-5252.1993

112. Manganelli R, Gennaro ML. Protecting from Envelope Stress: Variations on the Phage-Shock-Protein Theme. Trends Microbiol. 2017;25:205–16; doi:10.1016/j.tim.2016.10.001

113. Freeman ZN, Dorus S, Waterfield NR. The KdpD/KdpE Two-Component System: Integrating K+ Homeostasis and Virulence. PLoS Pathog. 2013;9:e1003201-; doi:10.1371/journal.ppat.1003201

114. Meadows JA, Wargo MJ. Carnitine in bacterial physiology and metabolism. Microbiology (N Y). 2015;161:1161–74; doi:10.1099/mic.0.000080

115. Kolata P, Efremov RG. Structure of *Escherichia coli* respiratory complex I reconstituted into lipid nanodiscs reveals an uncoupled conformation. Swartz KJ, Carter AP, Kumar V, Letts JA, editors. Elife. 2021;10:e68710; doi:10.7554/eLife.68710

116. Korea C-G, Badouraly R, Prevost M-C, Ghigo J-M, Beloin C. *Escherichia coli* K-12 possesses multiple cryptic but functional chaperone–usher fimbriae with distinct surface specificities. Environ Microbiol. 2010;12:1957–77; doi:10.1111/j.1462-2920.2010.02202.x

117. Fu H, Lee J, Wang T. Heat-shock increases rpoD-dependent β-galactosidase activity in the Escherichia coli strains BD792 and B23. 2015;19

118. Gagarinova A, Hosseinnia A, Rahmatbakhsh M, Istace Z, Phanse S, Moutaoufik MT, et al. Auxotrophic and prototrophic conditional genetic networks reveal the rewiring of transcription factors in *Escherichia coli*. Nat Commun. 2022;13:4085; doi:10.1038/s41467-022-31819-x

119. Prusiner S, Davis JN, Stadtman ER. Regulation of glutaminase B in *Escherichia coli*. I. Purification, properties, and cold lability. Journal of Biological Chemistry. 1976;251:3447–56; doi: 10;251(11):3447-56

120. Iacometti C, Marx K, Hönick M, Biletskaia V, Schulz-Mirbach H, Satanowski A, et al. Activating silent glycolysis bypasses *Escherichia coli*; bioRxiv. 2021;2021.11.18.468982; doi: 10.34133/2022/9859643

121. Njenga R, Boele J, Öztürk Y, Koch H-G. Coping with stress: How bacteria fine-tune protein synthesis and protein transport. Journal of Biological Chemistry. 2023;299:105163; doi:10.1016/j.jbc.2023.105163

122. Hajra D, Nair AV, Chakravortty D. An elegant nano-injection machinery for sabotaging the host: Role of Type III secretion system in virulence of different human and animal pathogenic bacteria. Phys Life Rev. 2021;38:25–54; doi:10.1016/j.plrev.2021.05.007

123. Peek J, Christendat D. The shikimate dehydrogenase family: Functional diversity within a conserved structural and mechanistic framework. Arch Biochem Biophys. 2015;566:85–99; doi:10.1016/j.abb.2014.12.006

124. Miwa T, Taguchi H. *Escherichia coli* small heat shock protein IbpA plays a role in regulating the heat shock response by controlling the translation of σ32. Proceedings of the National Academy of Sciences. 2023;120:e2304841120; doi:10.1073/pnas.2304841120

125. Brocklehurst KR, Hobman JL, Lawley B, Blank L, Marshall SJ, Brown NL, et al. ZntR is a Zn(II)-responsive MerR-like transcriptional regulator of zntA in *Escherichia coli*. Mol Microbiol. 1999;31:893–902; doi: 10.1046/j.1365-2958.1999.01229.x

126. T CP. A Galvanizing Story—Protein Stability and Zinc Homeostasis. J Bacteriol. 2007;189:2953–4; doi:10.1128/jb.00173-07

127. Iannuzzi C, Adrover M, Puglisi R, Yan R, Temussi PA, Pastore A. The role of zinc in the stability of the marginally stable IscU scaffold protein. Protein Sci. 2014;23(9):1208–1219. doi:10.1002/pro.2501.

128. Joly N, Engl C, Jovanovic G, Huvet M, Toni T, Sheng X, et al. Managing membrane stress: the phage shock protein (Psp) response, from molecular mechanisms to physiology. FEMS Microbiol Rev. 2010;34:797–827; 10.1111/j.1574-6976.2010.00240.x

129. LeRoux M, Culviner PH, Liu YJ, Littlehale ML, Laub MT. Stress Can Induce Transcription of Toxin-Antitoxin Systems without Activating Toxin. Mol Cell. 2020;79:280–292.e8; doi:10.1016/j.molcel.2020.05.028

130. Flores-Kim J, Darwin AJ. The Phage Shock Protein Response. Annu Rev Microbiol. 2016;70:83–101; doi:10.1146/annurev-micro-102215-095359

131. Walsh P, Bursać D, Law YC, Cyr D, Lithgow T. The J-protein family: modulating protein assembly, disassembly and translocation. EMBO Rep. 2004;5:567–571–571; doi:10.1038/sj.embor.7400172

132. Park DM, Akhtar MdS, Ansari AZ, Landick R, Kiley PJ. The Bacterial Response Regulator ArcA Uses a Diverse Binding Site Architecture to Regulate Carbon Oxidation Globally. PLoS Genet. 2013;9:e1003839; doi:10.1371/journal.pgen.1003839

133. Anand A, Patel A, Chen K, Olson CA, Phaneuf P V, Lamoureux C, et al. Laboratory evolution of synthetic electron transport system variants reveals a larger metabolic respiratory system and its plasticity. Nat Commun. 2022;13:3682; 10.1038/s41467-022-30877-5

134. Henry C, Loiseau L, Vergnes A, Vertommen D, Mérida-Floriano A, Chitteni-Pattu S, et al. Redox controls RecA protein activity via reversible oxidation of its methionine residues. Kowalczykowski SC, Storz G, editors. Elife. 2021;10:e63747; doi:10.7554/eLife.63747

135. Flint DH, Emptage MH, Guest JR. Fumarase A from *Escherichia coli:* purification and characterization as an iron-sulfur cluster containing enzyme. Biochemistry. 1992;31:10331–7; doi:10.1021/bi00157a022

136. Park S-J, Tseng C-P, Gunsalus RP. Regulation of succinate dehydrogenase (sdhCDAB) operon expression in *Escherichia coli* in response to carbon supply and anaerobiosis: role of ArcA and Fnr. Mol Microbiol. 1995;15:473–82; doi:10.1111/j.1365-2958.1995.tb02261.x

137. Roth M, Jaquet V, Lemeille S, Bonetti E-J, Cambet Y, Francois P, et al. Transcriptomic Analysis of *E. coli* after Exposure to a Sublethal Concentration of Hydrogen Peroxide Revealed a Coordinated Up-Regulation of the Cysteine Biosynthesis Pathway. Antioxidants. 2022;11; doi:10.3390/antiox11040655

138. Friedrich T, Dekovic DK, Burschel S. Assembly of the *Escherichia coli* NADH:ubiquinone oxidoreductase (respiratory complex I). Biochimica et Biophysica Acta (BBA) - Bioenergetics. 2016;1857:214–23; doi:10.1016/j.bbabio.2015.12.004

139. Prasad I, Young B, Schaefler S. Genetic Determination of the Constitutive Biosynthesis of Phospho-β-Glucosidase A in *Escherichia coli* K-12. J Bacteriol. 1973;114:909–15. 10.1128/jb.114.3.909-915.1973

140. Kuznetsova E, Proudfoot M, Gonzalez CF, Brown G, Omelchenko M V, Borozan I, et al. Genome-wide Analysis of Substrate Specificities of the *Escherichia coli* Haloacid Dehalogenase-like Phosphatase Family*. Journal of Biological Chemistry. 2006;281:36149– 61; doi:10.1074/jbc.m605449200

141. Sobota JM, Imlay JA. Iron enzyme ribulose-5-phosphate 3-epimerase in *Escherichia coli* is rapidly damaged by hydrogen peroxide but can be protected by manganese. Proceedings of the National Academy of Sciences. 2011;108:5402–7; doi:10.1073/pnas.1100410108

142. Chuanboon K, Na Nakorn P, Pannengpetch S, Laengsri V, Nuchnoi P, Isarankura-Na-Ayudhya C, et al. Proteomics and bioinformatics analysis reveal potential roles of cadmium-binding proteins in cadmium tolerance and accumulation of *Enterobacter cloacae*; PeerJ. 2019;7:e6904; https://pubmed.ncbi.nlm.nih.gov/31534833/

143. Imlay JA, Linn S. Bimodal pattern of killing of DNA-repair-defective or anoxically grown *Escherichia coli* by hydrogen peroxide. J Bacteriol. 1986;166:519–27; doi:10.1128/jb.166.2.519-527.1986

144. McCarthy DJ, Smyth GK. Testing significance relative to a fold-change threshold is a TREAT. Bioinformatics. 2009;25:765–71; doi:10.1093/bioinformatics/btp053

145. Umbarger HE, Brown B. Threonine deamination in *Escherichia coli*. II. Evidence for two L-threonine deaminases. J Bacteriol. 1957;73(1):105–12; doi:10.1128/jb.73.1.105-112.1957.

146. Julien M, Elisa G, Ronald VE, Enrique TMO, Ladan K, Laleh K, et al. Gene Erosion Can Lead to Gain-of-Function Alleles That Contribute to Bacterial Fitness. mBio. 2021;12:10.1128/mbio.01129-21; doi:10.1128/mbio.01129-21

147. Greenfield D, McEvoy AL, Shroff H, Crooks GE, Wingreen NS, Betzig E, et al. Self-Organization of the *Escherichia coli* Chemotaxis Network Imaged with Super-Resolution Light Microscopy. PLoS Biol. 2009;7:e1000137; doi:10.1371/journal.pbio.1000137

148. Guinote IB, Moreira RN, Barahona S, Freire P, Vicente M, Arraiano CM. Breaking through the stress barrier: the role of BolA in Gram-negative survival. World J Microbiol Biotechnol. 2014;30:2559–66; 10.1007/s11274-014-1702-4

149. Hemm MR, Paul BJ, Miranda-Ríos J, Zhang A, Soltanzad N, Storz G. Small stress response proteins in *Escherichia coli*: proteins missed by classical proteomic studies. J Bacteriol. 2010;192(1):46–58; doi:10.1128/JB.00872-09.

150. J RB, H DR, E PR, L SH, I BP, Aindrila M, et al. Functional Genomic Study of Exogenous n-Butanol Stress in *Escherichia coli*. Appl Environ Microbiol. 2010;76:1935–45; doi:10.1128/AEM.02323-09

151. Chen B, Chu T-W, Chiu K, Hong M-C, Wu T-M, Ma J-W, et al. Transcriptomic analysis elucidates the molecular processes associated with hydrogen peroxide-induced diapause termination in Artemia-encysted embryos. PLoS One. 2021;16:e0247160; doi:10.1371/journal.pone.0247160

152. Allison H, Jacquelyn S, Alexandra K, Kathleen C, Matthew Y, Dennis W, et al. The Bacterial iprA Gene Is Conserved across Enterobacteriaceae, Is Involved in Oxidative Stress Resistance, and Influences Gene Expression in *Salmonella enterica* Serovar Typhimurium. J Bacteriol. 2016;198:2166–79; doi:10.1128/jb.00144-16

153. Martínez Y, Li X, Liu G, Bin P, Yan W, Más D, et al. The role of methionine on metabolism, oxidative stress, and diseases. Amino Acids. 2017;49:2091–8; doi:10.1007/s00726-017-2494-2

154. Clausznitzer D, Oleksiuk O, Løvdok L, Sourjik V, Endres RG. Chemotactic Response and Adaptation Dynamics in *Escherichia coli*. PLoS Comput Biol. 2010;6:e1000784; doi:10.1371/journal.pcbi.1000784

155. Bouillet S, Hamdallah I, Majdalani N, Tripathi A, Gottesman S. A negative feedback loop is critical for recovery of RpoS after stress in Escherichia coli. bioRxiv. 2023;2023.11.09.566509; doi:10.1101/2023.11.09.566509.

156. Song S, Wood TK. A primary physiological role of toxin/antitoxin systems is phage inhibition. Front Microbiol. 2020;11:1895; doi:10.3389/fmicb.2020.01895.

157. chmidt M, Proctor T, Diao R, Freddolino PL. *Escherichia coli* YigI is a conserved Gammaproteobacterial acyl-CoA thioesterase permitting metabolism of unusual fatty acid substrates. J Bacteriol. 2022;204:e00014–22; doi:10.1128/JB.00014-22.

158. Jack RL, Sargent F, Berks BC, Sawers G, Palmer T. Constitutive expression of *Escherichia coli* tat genes indicates an important role for the twin-arginine translocase during aerobic and anaerobic growth. J Bacteriol. 2001;183(5):1801–4; doi:10.1128/jb.183.5.1801-1804.2001.

159. Zhao R, Li Z, Sun Y, Ge W, Wang M, Liu H, et al. Engineered *Escherichia coli* Nissle 1917 with urate oxidase and an oxygen-recycling system for hyperuricemia treatment. Gut Microbes. 2022;14:2070391; doi:10.1080/19490976.2022.2070391

160. Hryniewicz M, Sirko A, Pałucha A, Böck A, Hulanicka D. Sulfate and thiosulfate transport in *Escherichia coli* K-12: identification of a gene encoding a novel protein involved in thiosulfate binding. J Bacteriol. 1990;172(6):3358–66; doi:10.1128/jb.172.6.3358-3366.1990.

161. Sayers EW, Bolton EE, Brister JR, Canese K, Chan J, Comeau DC, et al. Database resources of the National Center for Biotechnology. 2022; doi:10.1093/nar/gkab1112

162. Krueger F, James F, Ewels P, Afyounian E, Weinstein M, Schuster-Boeckler B, et al. FelixKrueger/TrimGalore: v0.6.10 - add default decompression path. Zenodo; 2023; doi:10.5281/zenodo.7598955

163. FastQC. 2015. Available from: https://qubeshub.org/resources/fastqc

164. Langmead B, Salzberg SL. Fast gapped-read alignment with Bowtie 2. Nat Methods. 2012;9:357–9; doi:10.1038/nmeth.1923

165. Wang L, Wang S, Li W. RSeQC: quality control of RNA-seq experiments. Bioinformatics. 2012;28:2184–5; doi:10.1093/bioinformatics/bts356

166. Liao Y, Smyth GK, Shi W. featureCounts: an efficient general purpose program for assigning sequence reads to genomic features. Bioinformatics. 2014;30:923–30; doi:10.1093/bioinformatics/btt656

167. Ewels P, Magnusson M, Lundin S, Käller M. MultiQC: summarize analysis results for multiple tools and samples in a single report. Bioinformatics. 2016;32:3047–8. Available from: 10.1093/bioinformatics/btw354

168. Lamoureux CR, Decker KT, Sastry A V, Rychel K, Gao Y, McConn JL, et al. A multi-scale expression and regulation knowledge base for *Escherichia coli*. Nucleic Acids Res. 2023;51:10176–93; doi:10.1093/nar/gkad750

169. Consortium TGO, Aleksander SA, Balhoff J, Carbon S, Cherry JM, Drabkin HJ, et al. The Gene Ontology knowledgebase in 2023. Genetics. 2023;224:iyad031; doi:10.1093/genetics/iyad031

170. Gumienny R. GEOparse: Python Library to Access Gene ExpressionOmnibus Database(GEO). Available from: https://pypi.org/project/GEOparse/

171. Kim D, Seo SW, Gao Y, Nam H, Guzman GI, Cho B-K, et al. Systems assessment of transcriptional regulation on central carbon metabolism by Cra and CRP. Nucleic Acids Res. 2018;46:2901–17; doi:10.1093/nar/gky069

172. Spearman Rank Correlation Coefficient. The Concise Encyclopedia of Statistics. New York, NY: Springer New York; 2008. p. 502–5. Available from: 10.1007/978-0-387-32833-1_379

173. Martelli PL, Fariselli P, Casadio R. An ENSEMBLE machine learning approach for the prediction of all-alpha membrane proteins. Bioinformatics. 2003;19:i205–11. ; doi:10.1093/bioinformatics/btg1027

174. Pierleoni A, Martelli PL, Fariselli P, Casadio R. BaCelLo: a balanced subcellular localization predictor. Bioinformatics. 2006;22:e408–16; doi:10.1093/bioinformatics/btl222

175. Pierleoni A, Martelli PL, Casadio R. PredGPI: a GPI-anchor predictor. BMC Bioinformatics. 2008;9:392; doi:10.1186/1471-2105-9-392

176. Pierleoni A, Martelli PL, Casadio R. MemLoci: predicting subcellular localization of membrane proteins in eukaryotes. Bioinformatics. 2011;27:1224–30; doi:10.1093/bioinformatics/btr108

177. Savojardo C, Fariselli P, Casadio R. BETAWARE: a machine-learning tool to detect and predict transmembrane beta-barrel proteins in prokaryotes. Bioinformatics. 2013;29:504–5; doi:10.1093/bioinformatics/bts728

178. Szklarczyk D, Gable AL, Nastou KC, Lyon D, Kirsch R, Pyysalo S, et al. The STRING database in 2021: customizable protein–protein networks, and functional characterization of user-uploaded gene/measurement sets. Nucleic Acids Res. 2021;49:D605–12; 10.1093/nar/gkaa1074

179. Fernando PC, Mabee PM, Zeng E. Integration of anatomy ontology data with protein– protein interaction networks improves the candidate gene prediction accuracy for anatomical entities. BMC Bioinformatics. 2020;21:442; doi:10.1186/s12859-020-03773-2

180. Remmert M, Biegert A, Hauser A, Söding J. HHblits: lightning-fast iterative protein sequence searching by HMM-HMM alignment. Nat Methods. 2012;9:173–5; doi:10.1038/nmeth.1818

181. Bagheri H, Dyer R, Severin A, Rajan H. Comprehensive Analysis of Non Redundant Protein Database. 2020; doi:10.21203/rs.3.rs-54568/v1

182. Andreeva A, Kulesha E, Gough J, Murzin AG. The SCOP database in 2020: expanded classification of representative family and superfamily domains of known protein structures. Nucleic Acids Res. 2020;48:D376–82. Available from:10.1093/nar/gkz1064

183. Barrio-Hernandez I, Yeo J, Jänes J, Mirdita M, Gilchrist CLM, Wein T, et al. Clustering predicted structures at the scale of the known protein universe. Nature. 2023;622:637–45; doi:10.1038/s41586-023-06510-w

184. Tuttle AR, Trahan ND, Son MS. Growth and Maintenance of *Escherichia coli* Laboratory Strains. Curr Protoc. 2021;1:e20; doi:10.1002/cpz1.20

185. Schroeder A, Mueller O, Stocker S, Salowsky R, Leiber M, Gassmann M, et al. The RIN: an RNA integrity number for assigning integrity values to RNA measurements. BMC Mol Biol. 2006;7:3; doi:10.1186/1471-2199-7-3

186. Lataretu M, Hölzer M. RNAflow: An Effective and Simple RNA-Seq Differential Gene Expression Pipeline Using Nextflow. Genes (Basel). 2020;11:1487; doi:10.3390/genes11121487

187. Chen S, Zhou Y, Chen Y, Gu J. fastp: an ultra-fast all-in-one FASTQ preprocessor. Bioinformatics. 2018;34:i884–90; doi:10.1093/bioinformatics/bty560

188. Chen S, Zhou Y, Chen Y, Gu J. fastp: an ultra-fast all-in-one FASTQ preprocessor. Bioinformatics. 2018;34:i884–90; doi:10.1093/bioinformatics/bty560

189. Kim D, Langmead B, Salzberg SL. HISAT: a fast spliced aligner with low memory requirements. Nat Methods. 2015;12:357–60; doi:10.1038/nmeth.3317

190. Li H, Handsaker B, Wysoker A, Fennell T, Ruan J, Homer N, et al. The Sequence Alignment/Map format and SAMtools. Bioinformatics. 2009;25:2078–9.; doi:10.1093/bioinformatics/btp352

191. Liao Y, Smyth GK, Shi W. featureCounts: an efficient general purpose program for assigning sequence reads to genomic features. Bioinformatics. 2014;30:923–30; doi:10.1093/bioinformatics/btt656

192. Love MI, Huber W, Anders S. Moderated estimation of fold change and dispersion for RNA-seq data with DESeq2. Genome Biol. 2014;15:550; doi:10.1186/s13059-014-0550-8

193. Duda JC, Drenda C, Kästel H, Rahnenführer J, Kappenberg F. Benefit of using interaction effects for the analysis of high-dimensional time-response or dose-response data for two-group comparisons. Sci Rep. 2023;13:20804; doi:10.1038/s41598-023-47057-0

194. Liou G-G, Chao Kaberdina A, Wang W-S, Kaberdin VR, Lin-Chao S. Combined Transcriptomic and Proteomic Profiling of *E. coli* under Microaerobic versus Aerobic Conditions: The Multifaceted Roles of Noncoding Small RNAs and Oxygen-Dependent Sensing in Global Gene Expression Control. Int J Mol Sci. 2022;23; doi:10.3390/ijms23052570

