## Supplementary figures and table for "Annotating Hypothetical Proteins in *Escherichia coli* K-12"

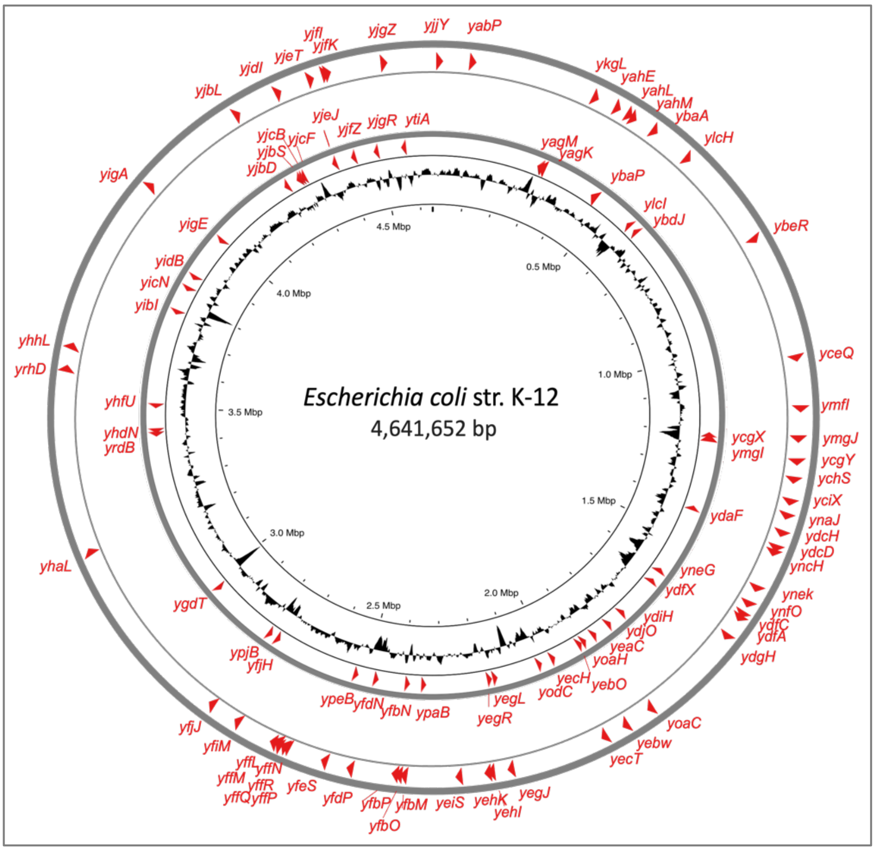


**Supplementary Figure 1: Circular representation of the *Escherichia coli* str. K-12 genome and distribution of hypothetical protein encoding genes**. Illustrated from the outside to the center of the circle: uncharacterized hypothetical protein encoding genes along with their strand orientations; GC content over a 10,000 bp window. Generated with Proksee [1].

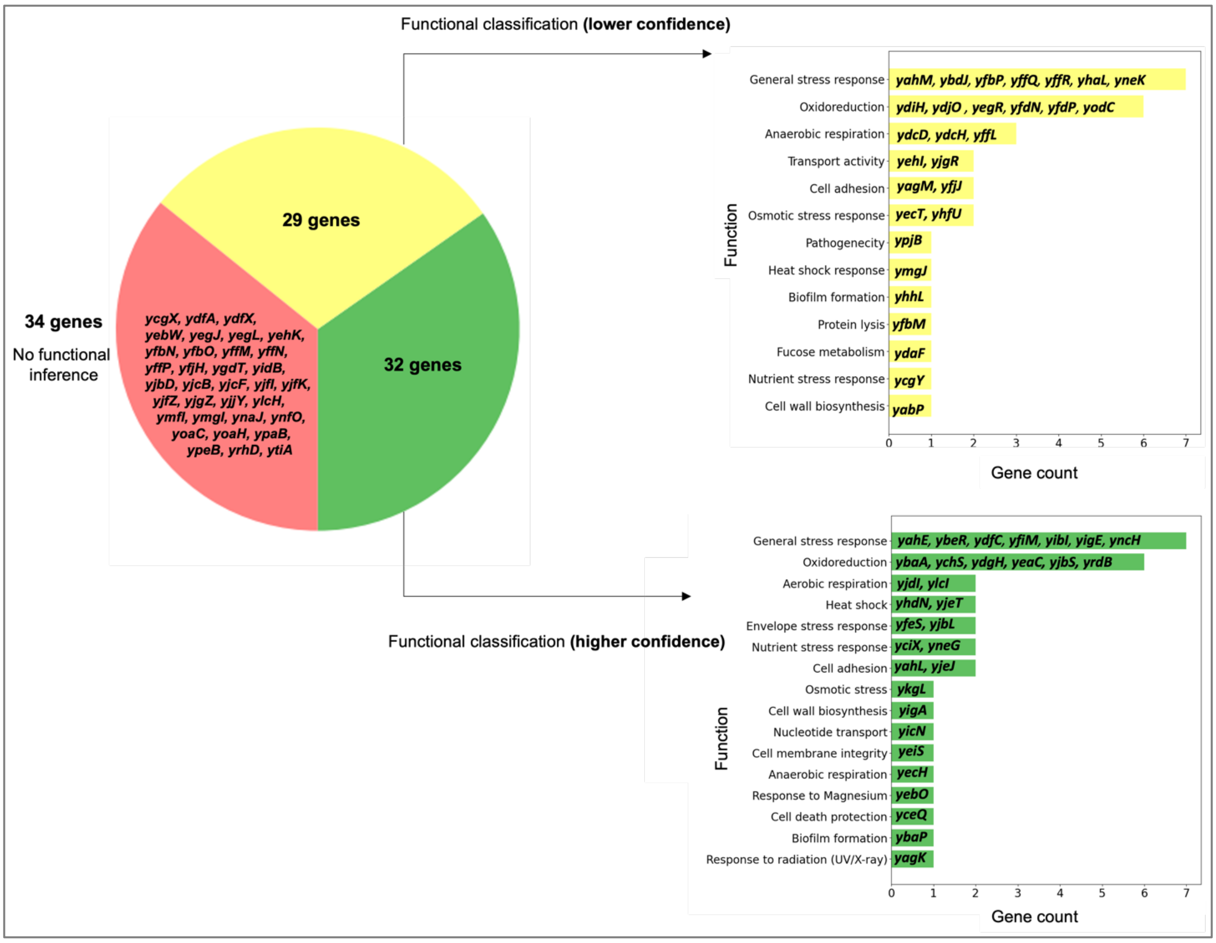


**Supplementary Figure 2: Classification of HP-encoding genes based on *in silico* tools regarding their confidence of assignment and functional categories.** 95 HP-encoding genes were classified into three categories. 32 genes (in green) could be assigned a function with a 'higher' confidence while 29 genes (yellow) were categorized with a 'lower' confidence. 34 genes (red) could not be functionally annotated. A 'higher' confidence implies well-correlated information from three or more *in silico* tools/databases (sources). A 'lower' confidence implies information could only be correlated from at least two sources. Genes with higher and lower confidence were functionally categorized based on all available *in silico* information (from manuscript Table 1.)

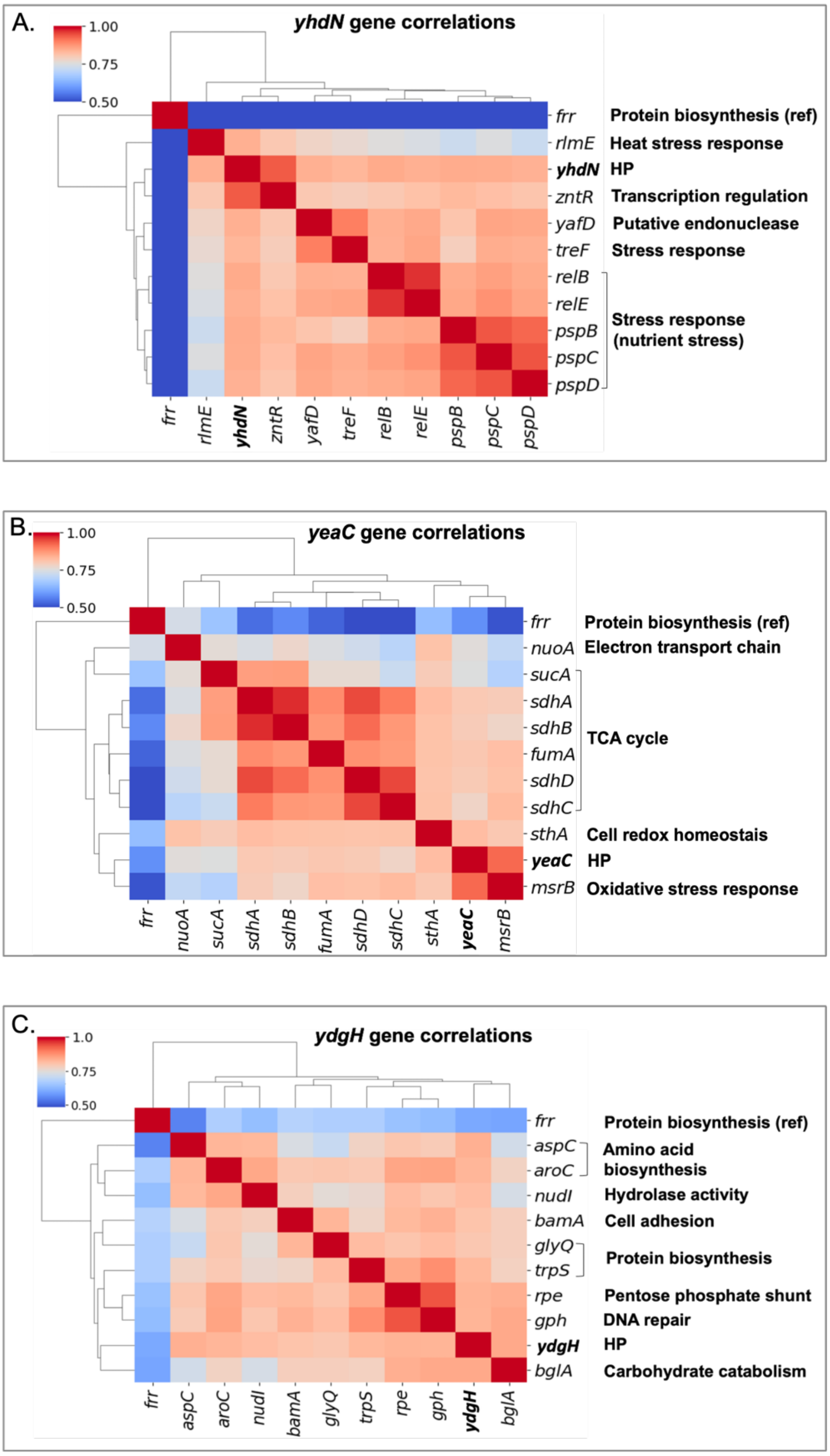


**Supplementary Figure 3: Gene co-expression analyses of three HP-encoding genes.** Heatmaps showing gene-gene correlations based on the Tjaden dataset [2]. The *frr* gene, encoding the ribosome – recycling factor, is included as a reference gene (see main text). Genes are clustered using the average linkage method based on their Euclidean distance, highlighting groups of genes with similar expression profiles, suggesting potential regulatory relationships or functional similarities. **A**. Top correlated genes with *yhdN,* **B**. *yeaC* and **C**. *ydgH*.

Abbreviations: HP, hypothetical protein

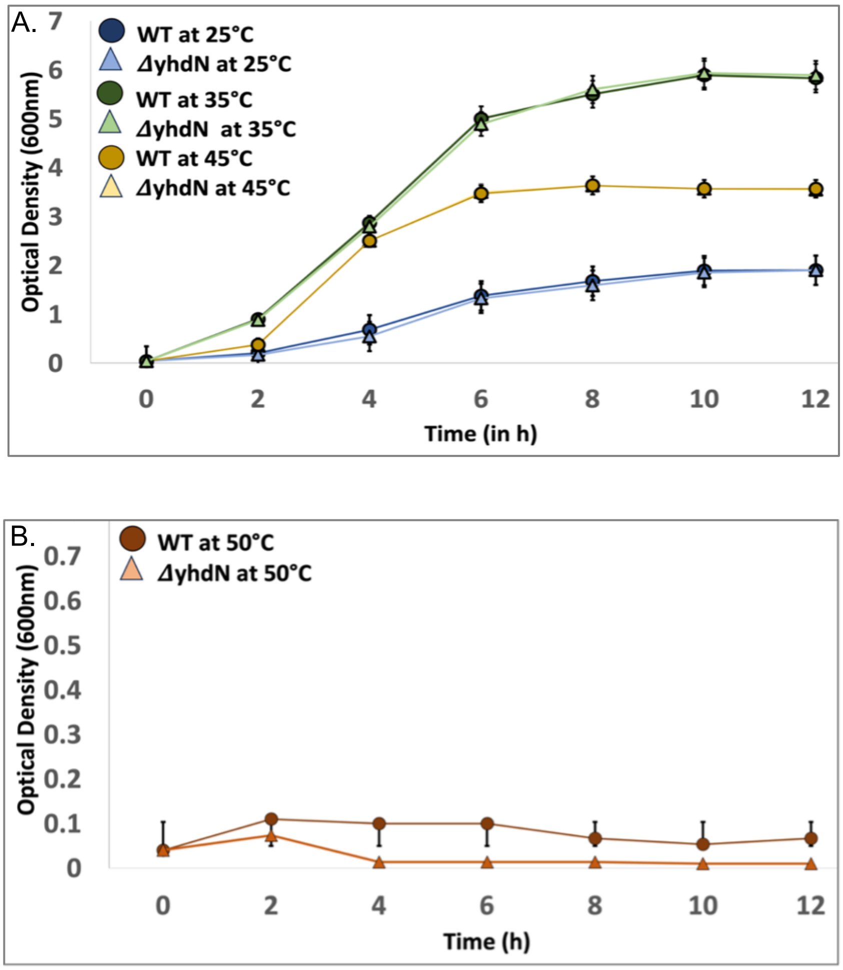


**Supplemental Figure 4: Growth curves of *E. coli* K-12 BW25113 wild type (WT) and *ΔyhdN* cells at different temperatures**. WT and respective mutant strain were continuously grown in LB medium for 12h at temperatures of **A.** 25°C, 35°C and 45°C for 12h (at starting $OD_{600}$ nm = 0.04); **B.** 50°C (at starting $OD_{600}$ nm = 0.04). The average of three independent readings was taken for each specified condition. Error bars on the graph indicate standard deviation from the mean.

### Suplementary Tables

**Supplementary Table 1.** Summary of HP-encoding genes associated with specific biological processes as derived from Protein ANalysis THrough Evolutionary Relationships (PANTHER) Gene Ontology (GO) categories [3]. Each gene is listed alongside its associated biological process, P-value, and false discovery rate (FDR). The P-values indicate the statistical significance of the association between the gene and the GO category, while the FDR values represent the adjusted probability of false positives. All obtained GO categories were statistically significant (*p*<0.05).

| Gene | PANTHER derived GO category | P-value | FDR |
|---|---|---|---|
| 1. *yabP* | Anaerobic respiration | 7.68E-05 | 4.53E-02 |
| 2. *yagM* | Cell adhesion | 1.49E-14 | 8.79E-12 |
| 3. *yagK* | Glycine betaine biosynthesis | 2.30E-09 | 2.26E-06 |
| 4. *yahE* | Not co-regulated | - | - |
| 5. *yahL* | Cell adhesion | 1.49E-14 | 8.79E-12 |
| 6. *yahM* | Glycine betaine biosynthesis | 2.30E-09 | 2.26E-06 |
| 7. *ybaA* | No functional group identified | - | - |
| 8. *ybaP* | No functional group identified | - | - |
| 9. *ybdJ* | Not co-regulated | - | - |
| 10. *ybeR* | Not co-regulated | - | - |
| 11. *yceQ* | Cell death | 7.79E-08 | 2.30E-04 |
| 12. *ycgX* | Not co-regulated | - | - |
| 13. *ycgY* | Ethanol/<br>Osmotic stress response | 2.24E-05<br>1.77E-05 | 9.43E-03<br>8.71E-03 |
| 14. *ychS* | No functional group identified | - | - |
| 15. *yciX* | No functional group identified | - | - |
| 16. *ydaF* | Fucose metabolic process | 2.49E-10 | 6.14E-08 |
| 17. *ydcD* | Phenylacetate catabolic process | 1.17E-15 | 3.47E-12 |
| 18. *ydcH* | Phenylacetate catabolic process | 1.17E-15 | 3.47E-12 |

| | | | |
|---|---|---|---|
| *19. ydfA* | Not co-regulated | - | - |
| *20. ydfC* | Not co-regulated | - | - |
| *21. ydfX* | Not co-regulated | - | - |
| *22. ydgH* | Oxidoreduction | 6.14E-05 | 2.00E-02 |
| *23. ydiH* | Fucose metabolism | 2.78E-06 | 9.04E-04 |
| *24. ydjO* | Response to acidic pH | 4.33E-04 | 4.73E-02 |
| *25. yeaC* | Aerobic respiration | 2.85E-13 | 4.21E-10 |
| *26. yebO* | Cellular response to magnesium ion | 4.21E-07 | 4.14E-04 |
| *27. yebW* | D-glucoronate catabolic process | 4.67E-05 | 6.90E-02 |
| *28. yecH* | Anaerobic respiration | 7.68E-05 | 4.53E-02 |
| *29. yecT* | Not co-regulated | - | - |
| *30. yegJ* | Not co-regulated | - | - |
| *31. yegL* | Not co-regulated | - | - |
| *32. yegR* | Oxalate catabolic process | | |
| *33. yehI* | Not co-regulated | - | - |
| *34. yehK* | Not co-regulated | - | - |
| *35. yeiS* | Not co-regulated | - | - |
| *36. yfbM* | Aerobic respiration | 2.85E-13 | 4.21E-10 |
| *37. yfbN* | Response to acidic pH | 4.33E-04 | 4.73E-02 |
| *38. yfbO* | Not co-regulated | - | - |
| *39. yfbP* | Not co-regulated | - | - |
| *40. yfdN* | Not co-regulated | - | - |
| *41. yfdP* | Not co-regulated | - | - |
| *42. yfeS* | No functional group identified | - | - |
| *43. yffL* | No functional group identified | - | - |
| *44. yffM* | Not co-regulated | - | - |
| *45. yffN* | Not co-regulated | - | - |
| *46. yffP* | Not co-regulated | - | - |
| *47. yffQ* | Not co-regulated | - | - |
| *48. yffR* | Not co-regulated | - | - |
| *49. yfiM* | Cell death | 7.79E-08 | 2.30E-04 |
| *50. yfjH* | Not co-regulated | - | - |
| *51. yfjJ* | Response to acidic pH | 4.33E-04 | 4.73E-02 |

| | | | |
|---|---|---|---|
| *52. ygdT* | Not co-regulated | - | - |
| *53. yhaL* | Not co-regulated | - | - |
| *54. yhdN* | Cellular response to heat | 5.24E-10 | 3.09E-07 |
| *55. yhfU* | Not co-regulated | - | - |
| *56. yhhL* | Not co-regulated | - | - |
| *57. yibI* | Response to nitrosative stress | 1.11E-05 | 3.29E-02 |
| *58. yicN* | Not co-regulated | - | - |
| *59. yidB* | Not co-regulated | - | - |
| *60. yigA* | Not co-regulated | - | - |
| *61. yigE* | Not co-regulated | - | - |
| *62. yjbD* | Not co-regulated | - | - |
| *63. yjbL* | Response to acidic pH | 4.33E-04 | 4.73E-02 |
| *64. yjbS* | Response to acidic pH | 4.33E-04 | 4.73E-02 |
| *65. yjcB* | Carbohydrate transport | 6.47E-05 | 1.74E-02 |
| *66. yjcF* | Response to acidic pH | 4.33E-04 | 4.73E-02 |
| *67. yjdI* | Oxidoreduction | 6.14E-05 | 2.00E-02 |
| *68. yjeJ* | Cell adhesion | 1.49E-14 | 8.79E-12 |
| *69. yjeT* | Not co-regulated | - | - |
| *70. yjfI* | Glycogen metabolic process | 1.72E-05 | 5.07E-02 |
| *71. yjfK* | Not co-regulated | - | - |
| *72. yjfZ* | Not co-regulated | - | - |
| *73. yjgR* | Not co-regulated | - | - |
| *74. yjgZ* | No functional group identified | - | - |
| *75. yjjY* | Cell death | 7.79E-08 | 2.30E-04 |
| *76. ykgL* | Glycine betaine biosynthesis | 2.30E-09 | 2.26E-06 |
| *77. ylcH* | Transcriptional attenuation by ribosome | 6.92E-06 | 2.92E-03 |
| *78. ylcI* | Anaerobic respiration | 7.68E-05 | 4.53E-02 |
| *79. ymfI* | DNA restriction/modification | 1.58E-09 | 4.67E-06 |
| *80. ymgI* | No functional group identified | - | - |

| | | | |
|---|---|---|---|
| *81. ymgJ* | No functional group identified | - | - |
| *82. ynaJ* | Aerobic respiration | 2.85E-13 | 4.21E-10 |
| *83. yncH* | Response to acidic pH | 4.33E-04 | 4.73E-02 |
| *84. yneG* | Not co-regulated | - | - |
| *85. yneK* | Not co-regulated | - | - |
| *86. ynfO* | | | |
| *87. yoaC* | Not co-regulated | - | - |
| *88. yoaH* | Anaerobic respiration | 7.68E-05 | 4.53E-02 |
| *89. yodC* | Anaerobic respiration | 7.68E-05 | 4.53E-02 |
| *90. ypaB* | Not co-regulated | - | - |
| *91. ypeB* | Oxidoreduction | 6.14E-05 | 2.00E-02 |
| *92. ypjB* | Not co-regulated | - | - |
| *93. yrdB* | Oxidoreduction | 6.14E-05 | 2.00E-02 |
| *94. yrhD* | Response to acidic pH | 4.33E-04 | 4.73E-02 |
| *95. ytiA* | Not co-regulated | - | - |

**References**

1. Grant JR, Enns E, Marinier E, Mandal A, Herman EK, Chen CY, et al. Proksee: in-depth characterization and visualization of bacterial genomes. Nucleic Acids Research. 2023 May 4;51(W1):W484–92. doi: 10.1093/nar/gkad326.
2. Tjaden B. Escherichia coli transcriptome assembly from a compendium of RNA-seq data sets. RNA Biol. 2023;20:77–84; doi:10.1080/15476286.2023.2189331
3. Thomas PD, Ebert D, Muruganujan A, Mushayahama T, Albou L-P, Mi H. PANTHER: Making genome-scale phylogenetics accessible to all. Protein Science. 2022;31:8–22; doi:10.1002/pro.4218
